# Mutational order and epistasis determine the consequences of *FBXW7* mutations during colorectal cancer evolution

**DOI:** 10.1101/2023.08.25.554836

**Authors:** Dedrick Kok Hong Chan, Amit Mandal, Yi Zhou, Scott David Collins, Richard Owen, James Bundred, Hannah Fuchs, Sabrina James, Iolanda Vendrell, Sarah Flannery, David Fawkner-Corbett, Jacob Househam, Trevor A Graham, Roman Fischer, Alison Simmons, Xin Lu, Simon James Alexander Buczacki

## Abstract

Somatic driver mutations, in genes such as *FBXW7,* have been discovered in phenotypically normal colonic tissue, however their role in cancer initiation remains elusive. Using normal and gene-edited patient-derived human colon organoids as models of early tumour evolution we observed that *FBXW7^-/-^* mutations exerted an epistatic effect on subsequent transcription depending on the mutational background of the cell. Specifically, the timing of acquiring an *FBXW7^-/-^* mutation respective to an *APC* mutation, led to profound phenotypic and transcriptomic differences. When *FBXW7* was mutated before *APC*, a near-normal cell state was maintained alongside repression of the *APC* transcriptional response. However, when *APC* was mutated before *FBXW7*, cells acquired classical cancer-stem cell features. Single-cell RNA sequencing revealed that mutation of *FBXW7* in normal tissue subtly switched cells from adult to a foetal/regenerative stem cell state. Further analysis using transposase-accessible chromatin sequencing identified this cellular plasticity was driven by changes in chromatin accessibility of transcriptional start site regions associated with TEAD, SNAI1 and AP-1 motifs, which in turn activate the foetal-like state. Taken together, we demonstrate a critical role of *FBXW7* mutations in preventing colorectal cancer initiation and provide exemplar evidence for the importance of epistasis and mutational order in cancer biology.

## Introduction

Somatic mutations have been detected in a range of normal human tissues, including the liver [1], oesophagus [2], skin [3], and endometrium [4]. Similarly, a catalogue of somatic mutations in normal human colonic epithelium using whole genome sequencing of two thousand crypts, strikingly found mutated cancer genes in phenotypically normal colonic epithelium [5]. In healthy colon such mutations form a distinct category from initiating cancer driving mutations in genes such as *TP53* or *APC*, which are ubiquitously found in colorectal cancer. *FBXW7* is a gene within the catalogue of mutations observed in normal colonic epithelium but its role in cancer initiation has not yet been investigated. FBXW7 is a key component of the Skp1-Cdc53/Cullin-F-box-protein complex (SCF/β-TrCP), an E3 ubiquitin ligase responsible for marking proteins by ubiquitination and targeting them for proteasomal degradation [6]. Substrates of FBXW7 are regulators of gene transcription, cell cycle progression or activators of signalling pathways, thus making *FBXW7* a pivotal tumour suppressor gene. Given *FBXW7* appears to acquire mutations before the widely accepted initiating mutations associated with colorectal cancer such as *APC*, *TP53*, *SMAD4* or *KRAS*, raises the possibility that mutated *FBXW7* could play a critical role in early cancer evolution. Furthermore, the timing of the *FBXW7* mutation suggests that the order of mutation acquisition may have important effects on phenotype.

Recently, there has also been significant interest in the effects of epigenetic plasticity in cancer evolution. In models of early pancreatic cancer, *KRAS*-mutant cells were shown to demonstrate distinct chromatin accessibility patterns which could predict divergence into either a benign or malignant fate [7]. In colorectal cancer, the impact of epigenetic changes on tumour initiation are beginning to be uncovered. Johnstone et al. highlighted the importance of the three-dimensional architecture, in the form of chromatin loops, topologically activating domains and large-scale compartments, of DNA in restricting malignant progression [8]. A more recent study by Heide et al. found chromatin modifier genes are actively mutated in the early initiation of colorectal cancer and can result in the accumulation of further genetic mutations which herald carcinogenesis [9].

In this study, we used normal human adult and foetal colonic organoids, applying CRISPR-Cas9 gene editing to explore the cell autonomous and gene-interactive effects of an *FBXW7* mutation. Combining gene editing with organoid culture fashions a powerful tool ideal for studying the functional consequences of driver mutations in normal tissue for it allows specific mutations to be introduced into tissue devoid of genetic perturbation. By sequentially introducing multiple distinct mutations into the same cell, we also aimed to determine relationships arising between cells with mutant *FBXW7* and the acquisition of subsequent cancer driver mutations. We found that cells with mutant *FBXW7* were transcriptionally like wildtype cells but primed towards a foetal stem cell state. Further, we observed that an initial *FBXW7* mutation repressed the transcriptional effect of a subsequent *APC* mutation, resulting in retention of the near-wildtype phenotype. Whereas when we reversed this order of mutations, despite identical genotype, we generated cancer stem cell organoids. We uncovered this profound mutational order-based cellular plasticity to be driven by epigenetic differences in both global and local chromatin accessibility. Here we provide direct evidence that epistasis, and the order of mutations, as much as the mutations themselves, are key in determining phenotypes.

## Results

### Mutant *FBXW7^-/-^* recapitulate known effects on downstream targets

Patient-derived adult stem cell wildtype organoids (W) were generated from normal colon tissue of patients undergoing surgery for colorectal cancer. CRISPR-Cas9 editing was used to generate *FBXW7^-/-^* organoids (F) [10]. Our gene editing strategy utilizes three sgRNAs to introduce large deletions at exon 2 of the *FBXW7* gene. F and W organoids possessed similar morphology (Fig. 1a). We validated knockout of the *FBXW7* gene using western blot (Fig. 1b), and immunofluorescence (Fig. 1c). Western blotting of known FBXW7 targets, found upregulation of the expected substrates upon loss of FBXW7 (Fig. 1d).

**Fig. 1.**
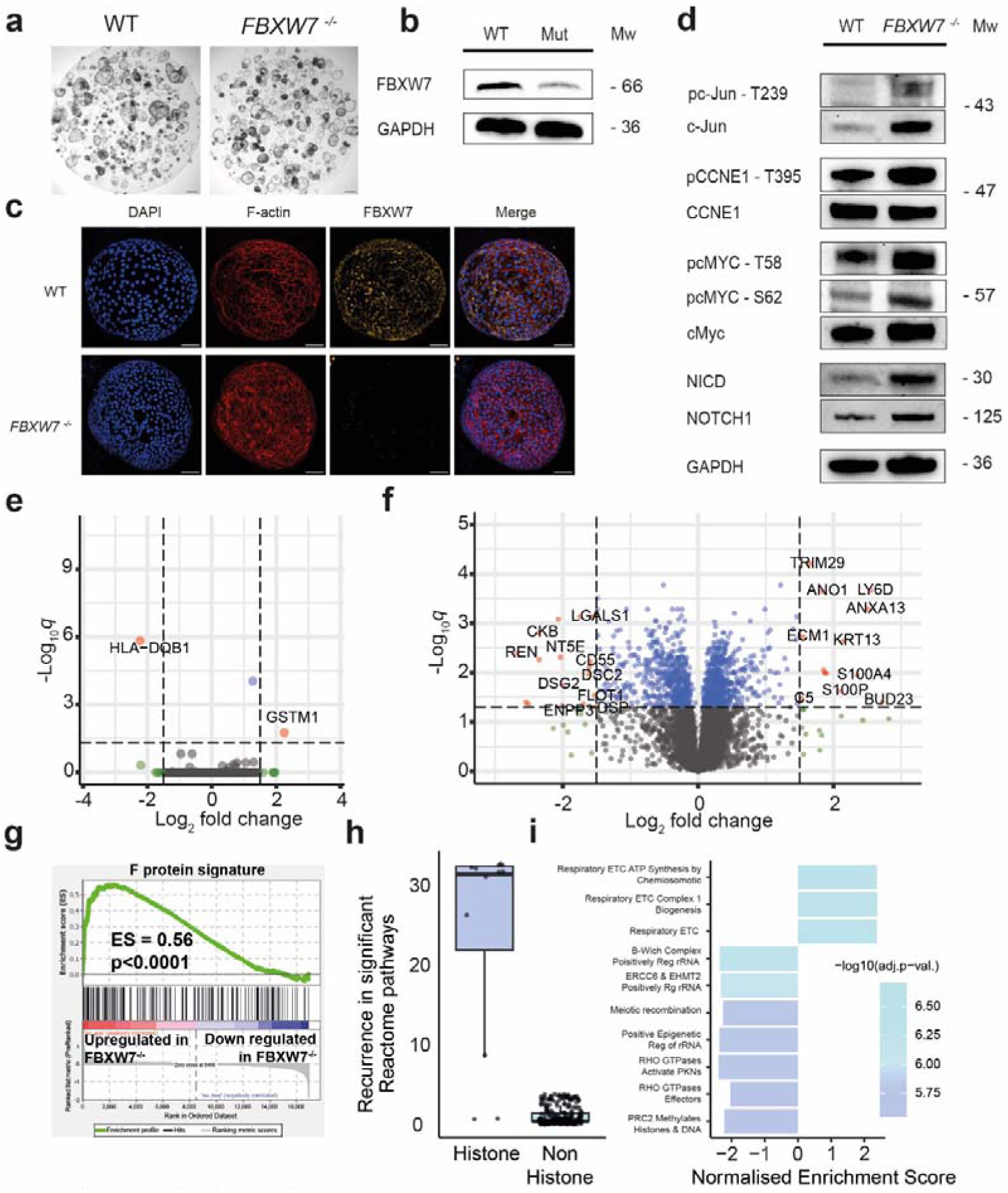
Characterisation of W and F organoids. **a,** Brightfield microscopy of W and F organoids revealed no observable phenotypic differences. Scale bars represent 500µm. **b.** Western blot validation showing loss of FBXW? in F organoids. **c.** lmmunofluorescence of Wand F organoids with DAPI nuclear stain (blue), F-actin (red), FBXW7 (orange). Scale bars represent 100µm. **d**, Western blot analysis of known ubqiutiination substrates of FBXW?. **e.** Volcano plot of bulk RNAseq of F vs W organoids revealed minimal differential expression of genes. Positive fold change = up in F. **f**, Volcano plot of proteomic MS analysis of F and W organoids. Positive fold change= up in F. **g**, GSEA comparing the top 200 proteins over expressed in F compared to the ranked gene list of differentially expressed transcripts between F and W organoids. **h**, Box and whisker plot for the recurrence of histone pathways in depleted Reactome pathways in the F vs W proteomics. Median+/− IQR. **i**, Chart detailing the enrichment scores for the top 10 Reactome pathways altered In F vs W proteomics. All experiments were performed with n=3 biological replicates (proteomics n=4).

To characterise transcriptional differences between F and W organoids, we performed bulk-RNA sequencing (bulk-RNAseq) (Fig. 1e). Using a cut-off of log_2_FC ≤ −1.5 or ≥ 1.5, and adjusted *p*-value < 0.05, F and W were transcriptionally similar with only two genes found differentially expressed. Proteomic comparison of F and W organoids showed a larger number of differentially expressed proteins (n=1097, adjusted *p*-value < 0.05) however most of those identified showed only small fold changes. When applying the same fold change cut-off as used in bulk-RNAseq, this dropped to 31 proteins (log_2_FC ≤ −1.5 or ≥ 1.5, and adjusted *p*-value < 0.05) (Fig. 1f). Gene set enrichment analysis (GSEA) of the top 200 upregulated proteins in F organoids and the ranked bulk-RNAseq gene list comparing F and W organoids showed strong enrichment (Fig. 1g). Pathway analysis of the ranked proteins revealed 42 significant Reactome pathways (5 enriched and 37 depleted: adjusted p-value < 0.01 and NES ≤ −2 or ≥ 2). Interestingly, histone-related proteins contributed significantly (Mann-Whitney p-value = 1.64e-07) to the depleted Reactome pathways (Fig. 1h,i). The average recurrence of proteins H2AC21, H2AC6, H2AC7, H2AX, H2AZ1, H2BC11, H2BC17, H2BC26, H2BC3, H2BC5, H3-3A and H4C1 was 24.9, whereas it was 1.7 only for the remaining set of total 270 proteins contributing to the leading-edge of the enriched/ depleted pathways (Extended Data Table 1). In summary these analyses confirm *FBXW7^-/-^* mutations on a wildtype background have a subtle effect on phenotype and transcriptome although proteomic changes were apparent.

### *FBXW7^-/-^* mutants do not exhibit a competitive proliferative advantage over wildtype organoids

Given the transcriptional similarities between W and F organoids, we interrogated the possibility for competitive or cooperative effects by co-culturing F organoids with W organoids, analogous to the recently described supercompetitor effect seen with *Apc^-/-^*intestinal organoids [11]. *APC^-/-^* (A) organoids were also generated by targetting the hotspot region using CRISPR-Cas9 editing to introducing a biallelic frameshift indel at codon 1499. A were co-cultured with W organoids and A were found to have a significant proliferative advantage in concordance with previous murine data (Extended Data Fig. 1a,c,d). F organoids however did not demonstrate any advantage when co-cultured with W organoids and transcriptional profiling showed minimal differences between W:W-cocultured and F:Fcocultured organoids (Extended Data Fig. 1b, e-i) in contrast to our previous findings in colorectal cancer cell lines [12].

### *FBXW7* mutation provides evidence of epistasis

In light of the lack of a cooperative/competitive interaction with wildtype cells and building on our earlier finding that F organoids were transcriptionally similar to W organoids, we generated an organoid model mirroring the polypoidal and cancerous phases of CRC carcinogenesis (Fig. 2a). A organoids represent the polypoidal, precancerous phase, while a *TP53^-/-^* mutation introduced into A organoids, making a double knockout *APC^-/-^ / TP53^-/-^* mutation, generated AT organoids, representing the cancerous phase. Successful targeting was confirmed using Sanger sequencing and western blotting (Extended Data Fig. 2a). On each of these A and AT organoids, an *FBXW7* mutation was then introduced, generating AF and ATF organoids respectively (Fig. 2b) (Extended Data Table 2).

**Fig. 2.**
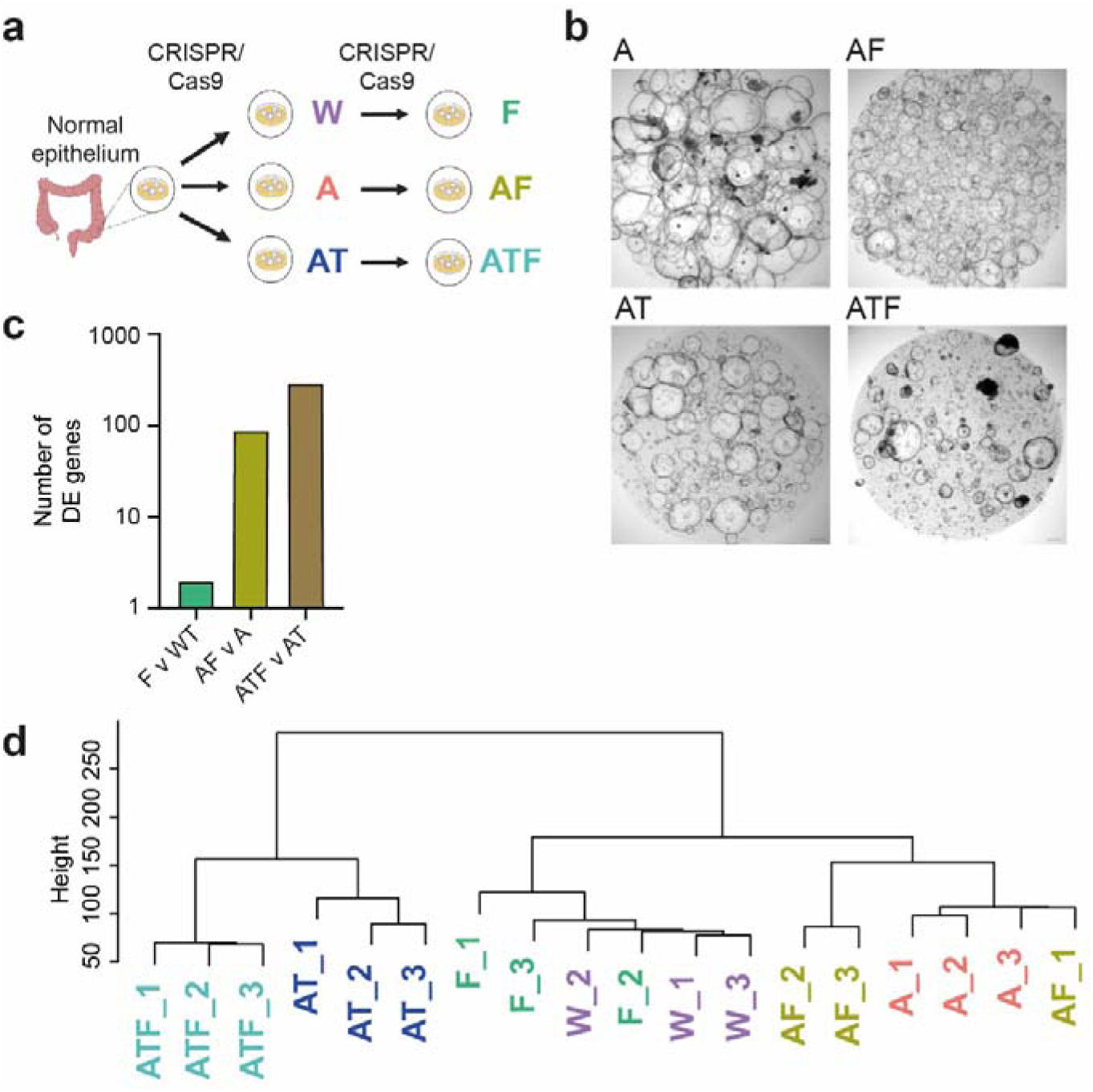
Effect of FBXW7 mutation is dependent on mutational background. **a,** Schematic showing the organoid models used. W, A and AT organoids were generated to reflect normal, adenomatous and carcinomatous lesions. On each of these organoid types, an FBXW7 mutation was introduced to generate F, AF and ATF organoids respectively. **b,** Brightfield microscopy of A, AF, AT and ATF organoids. A organoids adopt a cystic phenotype which is representative of increased proliferation. AF organoids appear smaller in size compared with A organoids. AT organoids also recapitulate the cystic appearance of A organoids. ATF organoids appear less cystic than AT organoids. Scale bars represent 100µm. **c,** Histogram showing the number of differentially expressed genes on the addition of an *FBXW7* mutation changes based on the mutational background. While 2 genes were differentially expressed between Wand F, 87 and 294 genes were differentially expressed between A and AF, and AT and ATF respectively. **d,** Hierarchical clustering, based on gene expression data, of the different organoid types showed that Wand F organoids clustered most similarly together. All experiments were performed with n=3 biological replicates.

As previously described, A organoids were morphologically distinct from W organoids, growing large and cystic [13, 14]. The subsequent acquisition of *TP53* mutation preserved this phenotype. Importantly, the additional mutation of *FBXW7* on both the A and AT organoids also maintained this cystic morphology. To further characterise transcriptional differences between these different organoids bulk-RNAseq of A organoids were compared with AF, while AT organoids were compared with ATF organoids. A organoids were compared with AF organoids, while AT organoids were compared with ATF organoids. Together with our earlier comparison between F and W organoids which found only 2 differentially expressed genes, we observed 87 differentially expressed genes between A and AF, and 294 between AT and ATF (Fig. 2c, and Extended Data Fig. 2b and 2c). Hierarchical clustering of these different groups showed that F and W organoids clustered together, while A, AF and AT, ATF organoids formed separate clusters (Fig. 2d). These results demonstrate an epistatic effect with the loss of *FBXW7* having varying effects on the cells depending on the mutational background. More importantly, it suggests that an *FBXW7* mutation can influence different cellular phenotypes depending on when the mutation was acquired, and that an early mutation will have fewer effects compared with a later mutation.

To ensure our observations were not an artefact of repeated iterations of gene editing, we targeted the *AAVS1* safe harbour locus three separate times to replicate the triple-mutant ATF organoids. Morphologically, *AAVS1*-targetted organoids resembled wildtype organoids (Extended Data Fig. 3a). Bulk-RNAseq of *AAVS1*-targetted organoids compared with W organoids showed only two differentially expressed genes with a cut-off of log_2_FC ≤ −1.5 or ≥ 1.5, and adjusted *p*-value < 0.05 (Extended Data Fig. 3b), greatly differing from the 493 differentially expressed genes between ATF and W organoids (Extended Data Fig. 3c). These findings confirm that the epistatic effects described earlier were not simply a function of repeated iterations of CRISPR-Cas9 gene editing.

### Order of *FBXW7* mutation determines effect on transcriptional profile

To ascertain the transcriptional effect of order of mutation, we designed an experiment in which organoids possessed the same mutations, but differed in the order in which the mutations were introduced. As previously described, AF organoids possessed *APC^-/-^* followed by *FBXW7^-/-^*. We further generated FA organoids, which acquired *FBXW7^-/-^*mutation prior to the *APC^-/-^* (Fig. 3a). Morphologically, FA organoids did not acquire the cystic phenotype characteristic of A or AF organoids (Fig. 3b). Principal component analysis (PCA) of AF and FA organoids processed for bulk-RNAseq, revealed that together with W, A and F organoids gene expression of FA organoids did not cluster with AF organoids (Fig. 3c). Instead, FA organoids clustered with F and W organoids, while AF organoids clustered with A organoids. Organoid clustering for all samples was independent of patient donor (Extended Data Fig. 3d). To analyse the transcriptional effect mutational order, we used a cut-off of log_2_FC ≤ −0.7 or ≥ 0.7 to identify differentially expressed genes when AF and FA organoids groups were compared with W organoids. This cut-off selected for 3335 genes in the AF group and 861 genes in the FA group (Fig. 3d). There was a significant overlap of the FA genes where 628 (72.9%) were also differentially expressed in the AF organoids compared to W. Further analyses of the gene expression changes for these 628 genes suggested a repressive effect after an initial *FBXW7* mutation, such that the magnitude of the transcriptional perturbation was rendered less positive or negative (Fig. 3e). Using GSEA, we also observed that some key signalling pathways such as the EMT pathway were significantly more prevalent in FA compared with AF organoids (normalised enrichment score (NES) 2.21, adjusted *p*-value= 1.70 × 10^-10^). Collectively, these findings promulgate our observation that mutational background alters the transcriptional effect of a mutation. Moreover, our findings also suggest that early loss of *FBXW7* in normal tissue may protect against the future effect of *APC* loss through transcriptional repression, and by extension could be protective against the initiation of CRC.

**Fig. 3.**
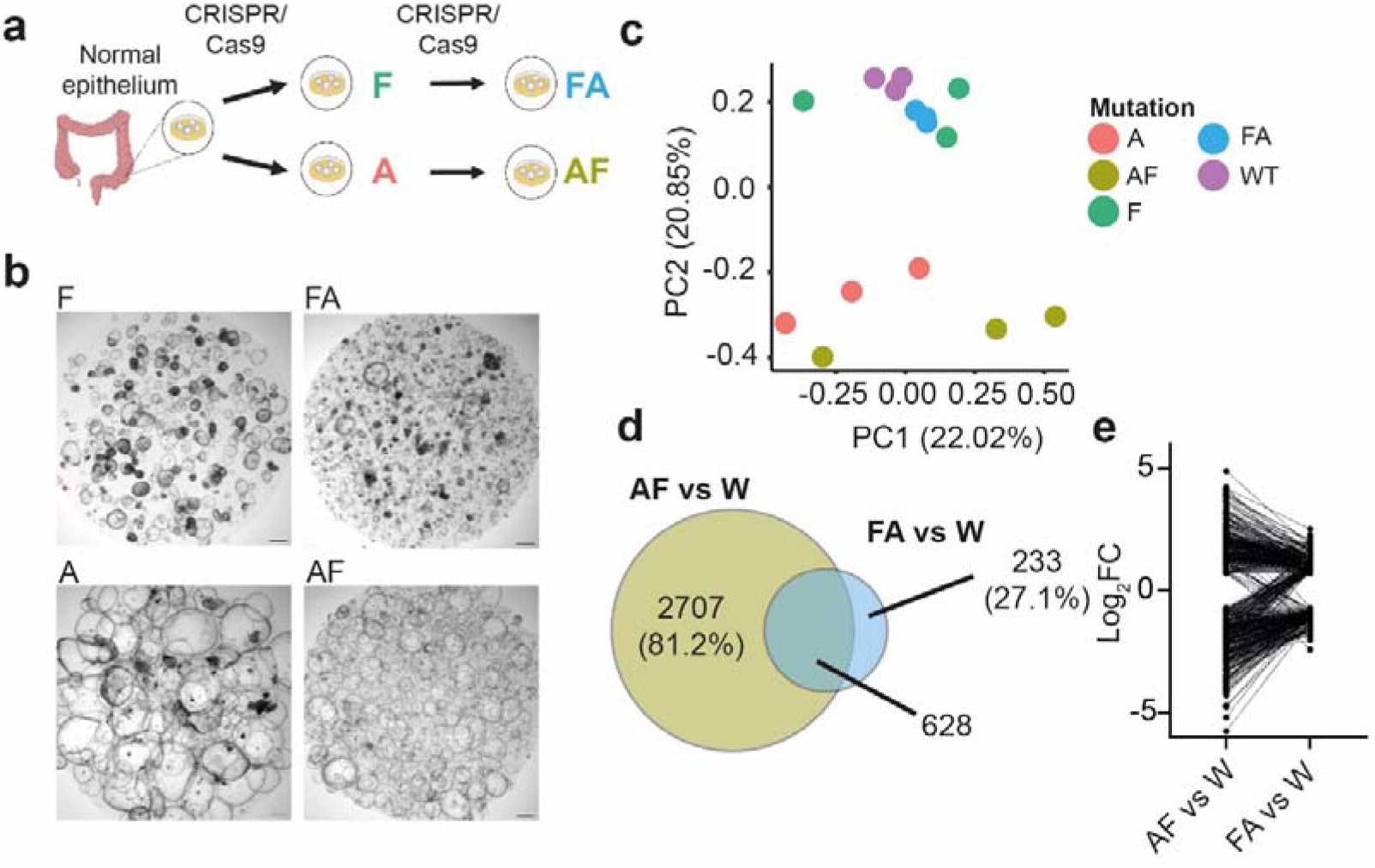
Effect of FBXW7 mutation is dependent on the order of mutation. **a,** Schematic showing the organoid models used in this analysis. A and F organoids were generated separately. On the A organoid, an *FBXWl* mutation was introduced, generating AF organoids. On the F organoid, an *APC* mutation was introduced, generating FA organoids. AF and FA organoids possessed the same mutations but acquired the mutations in an inverse order. **b,** Brighlfield microscopy *of* A, AF, F and FA organoids. Although FA organoids acquired an *APC* mutation, this did not recapitulate the cystic phenotype observed in A or AF organoids. Scale bars represent 100µm. **c,** Principal component analysis (PCA) showed not only that FA organoids clustered separately from AF organoids, FA organoids clustered together with W and F organoids, while AF organoids clustered closer to A organoids. **d,** Venn diagram depicting the overlap of differentially expressed genes at cut-off log2FC ≤ −0.7 or ≥ 0.7 between AF vs Wand FA vs W comparisons. 628 genes were differentially expressed in both AF and FA organoids. **e,** Linked column graph of AFvW and FAvW overlap genes (n=628) showed there was a reduction in the magnitude by which the genes were differentially expressed. All experiments were performed with n=3 biological replicates.

### scRNAseq data shows de-differentiation towards a foetal state in *FBXW7* mutant organoids

We endeavoured to explain our finding where an early *FBXW7* mutation was protective to consequences of a later acquisition of an *APC* mutation, considering our observation that mutational order affects transcription. We hypothesised that subtle transcriptomic changes could be present in F organoids, in small subpopulations of cells, which might be masked by bulking during RNAseq. To overcome this, we performed single-cell RNA sequencing (scRNAseq) on F and W organoids. Differential gene expression analysis using pseudobulked F and W cells showed a significant enrichment of upregulated F specific genes between the scRNAseq samples and our previously generated F and W bulk-RNAseq (p<0.005) (Extended Data Fig. 4a). Clustering by Uniform Manifold Approximation and Projection (UMAP) uncovered 14 clusters (Fig. 4a). Notably cluster 13 was unique to *FBXW7* mutant cells (Fig. 4b). This cluster was marked by genes such as *TAGLN* (log_2_FC= −1.33, *p*= 2.81 × 10^-35^) and *MMP7* (log_2_FC= −1.47, *p*= 7.70 × 10^-34^) and possessed an EMT signature analogous to that recently demonstrated in a separate analysis of murine foetal organoids (Extended Data Fig. 4b) [15]. Importantly, we observed that W cells expressed *FBXW7*, indicating that this gene is expressed in normal epithelium (Fig. 4c). Analysis by RNA velocity indicated that cluster 13 was an important terminal differentiation state, which could only have occurred in *FBXW7* mutant cells (Fig. 4d). As expected, quantitative analysis between UMAP plots from F and W organoids demonstrated a statistically significant upregulation of the FA_up gene signature in F organoids relative to W organoids (Fig. 4e). We also observed increased expression of the foetal gene signature [16] (Fig. 4f), and the YAP pathway gene signature [17] (Fig. 4g) in F organoids (Extended Data Table 3). There was also a decreased expression of the adult intestinal stem cell signature (ISC) [18] (Fig. 4h), and decreased expression of the adult crypt-based columnar (CBC) stem cell signature [19] (Fig. 4i) in F organoids. Reanalysis of our bulk-RNAseq data comparing F and W organoids using GSEA found that foetal (Extended Data Fig 5a) and YAP (Extended Data Fig. 5b) signatures correlated with our scRNAseq results. Similarly, overlaying our bulk-RNAseq F signature on a previously published scRNAseq analysis of foetal and adult colonic epithelia demonstrated expression of F-specific genes in foetal cells and W specific genes such as *OLFM4* in adult cells (Extended Data Fig. 5c) [20]. Immunofluorescence staining for LY6D validated it as a novel foetal marker; having been found consistently upregulated in F using bulk bulk-RNAseq, scRNAseq and proteomics. (Extended Data Fig. 5d).

**Fig. 4.**
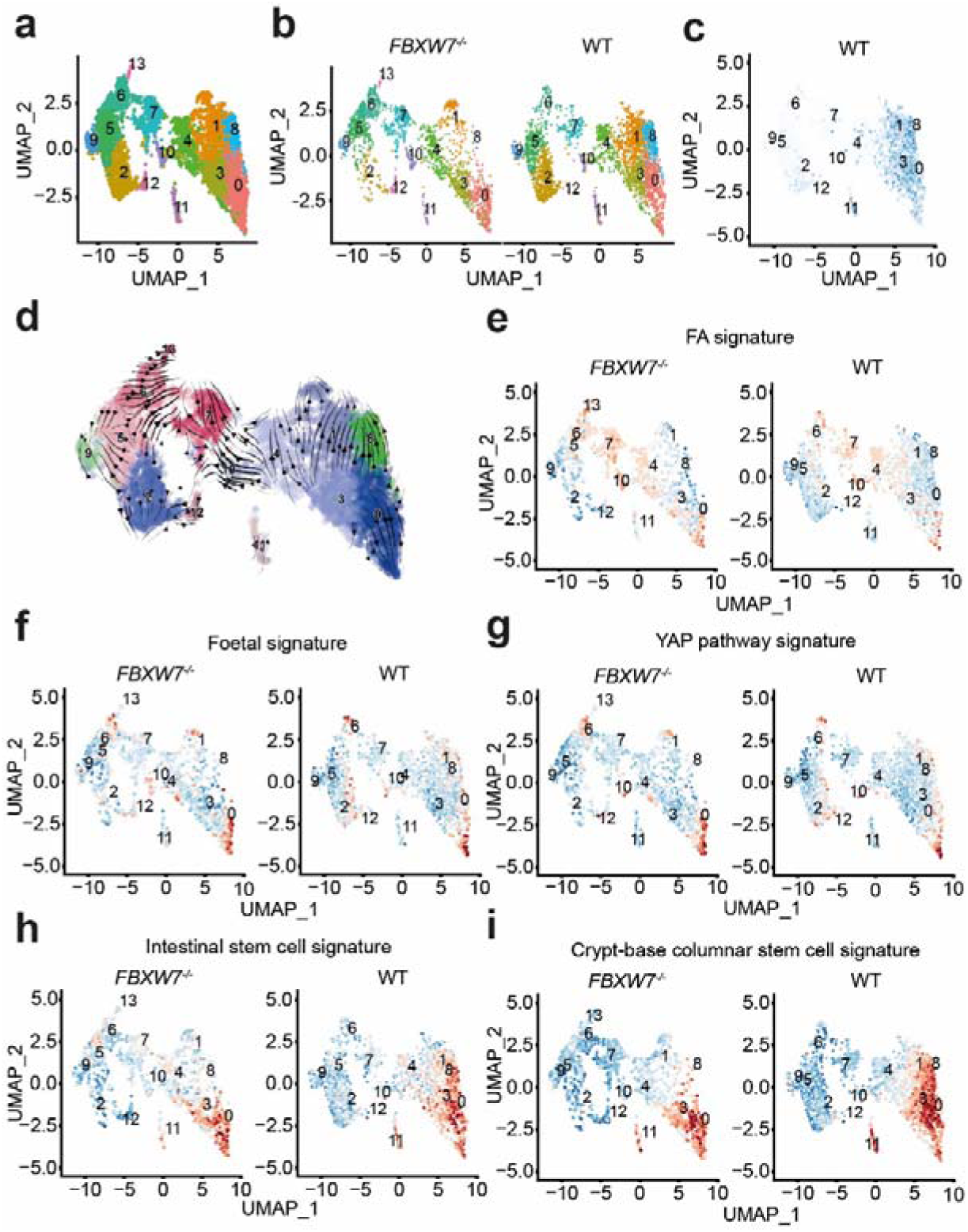
scRNAseq shows FBXW7 mutation induces a switch from an adult to foetal stem cell state. **a,** Uniform manifold approximation and projection (UMAP) analysis from scRNAseq of W and F organoids yielded 14 clusters. **b,** Cluster 13 was unique to F organoids. **c,** *FBXW7* is actively expressed in wildtype organoids. Intensity of blue reflects *FBXW7* level. **d,** RNA velocity analysis demonstrated that clusters 1, 8 and 13 were terminally differentiating subpopulations. Chi-square analysis of Wand F organoids demonstrated that F organoids were enriched for **(e)** FA_up gene signature (p<2.22 × 10·^16^), **(f)** foetal signature (p=9.4754 × 10 ^11^), **(g)** YAP pathway signature (p= 3.2134 × 10-^7^), and depleted for **(h)** intestinal stem cell signature (p=1.4887 × 10-6) and **(i)** crypt-based columnar stem cell signature (p<2.22 × 10 ^16^) Red *=* increased expression, Blue *=* decreased expresion. Data presented represents two biological replicates for each condition (n=4).

In F vs W organoids we also observed a downregulation of the adult-ISC (Extended Data Fig. 5e) and adult-CBC (Extended Data Fig. 5f) signatures in bulk-RNAseq as in scRNAseq, though this did not achieve statistical significance. To further explore the YAP pathway, we performed additional GSEA on FA vs W and AF vs W organoids. Intriguingly, this showed greater enrichment of the YAP signature in FA vs W (ES= 0.58) (Extended Data Fig. 6a) compared to AF vs W (ES= 0.36) (Extended Data Fig. 6b). Taken together, these findings point towards a plasticity in stem cell identity, favouring pathways which recall a foetal regenerative/YAP active stem cell state following *FBXW7* mutation.

In contrast to F organoids acquiring a foetal stem cell state, bulk-RNAseq comparing A vs W showed loss of the adult ISC signature and acquisition of the recently described proliferative colon stem cell (proCSC) signature in line with findings of others [21] (Extended Data Fig 7a,b). Further comparison of FA and AF transcriptomes showed that AF organoids retained a greater enrichment for the proCSC state whereas FA organoids developed even stronger enrichment for the foetal stem cell state than seen with the initial F mutation (NES Foetal: FA vs W = 2.03; F vs W= 1.99, p<0.0001) (Extended Data Fig 7c and d). To better dissect stem cell changes associated with acquisition of an *APC* mutation we performed scRNAseq on W and A organoids. scRNAseq recapitulated bulk-RNAseq findings demonstrating clear plasticity in stem cell populations with the proCSC population absent in W organoids but a dominant population in A organoids (Extended Data Fig 7e). In summary these data demonstrate that stem cell identity can be determined by order of mutations.

### Transcriptional effects of mutational order are driven by local and global changes in chromatin accessibility

We next sought to uncover the mechanism which resulted in FA and AF organoids exhibiting such profound transcriptional differences despite their identical genotypes. We hypothesised that changes in chromatin accessibility induced by a preceding *FBXW7* mutation could influence the effects of a subsequent *APC* mutation. To interrogate this, we performed assay for transposase-accessible chromatin with sequencing (ATACseq) on W, F, A, FA, AF and human foetal (Fo) colon organoids (17pcw and 21pcw), as an *in vitro* epigenomic model of tumour evolution. Initially we compared W, F, A (matched for donor) and foetal organoids finding whole genome accessibility to be greater in F, A and foetal organoids when compared to W (Fig. 5a). Foetal and F organoids possessed most euchromatin, compatible with the profound loss of histones seen in our proteomic analysis. We performed a *de novo* transcription factor motif scan using the GADEM algorithm on ATACseq peaks for all comparisons. Accessibility to TEAD, AP-1 and SNAI1 motifs, previously shown to be associated with the foetal/YAP state, were highly enriched in both foetal and F organoids (Extended Data Table 4) (Fig. 5b and c) [15]. However, the genomic loci of these motif sites were distinct between foetal and F organoids suggesting epigenetic differences between induced oncogenic (onco-foetal) and intrinsic foetal states. As expected, the F vs WT and FA vs AF transcriptional signatures were also found more accessible in F and foetal organoids (Extended Data Fig. 8a and b). In line with our bulk-RNAseq, scRNAseq and proetomics data we also found accessibility to *LY6D* to be greater in F organoids (Extended Data Fig. 8c).

**Fig. 5.**
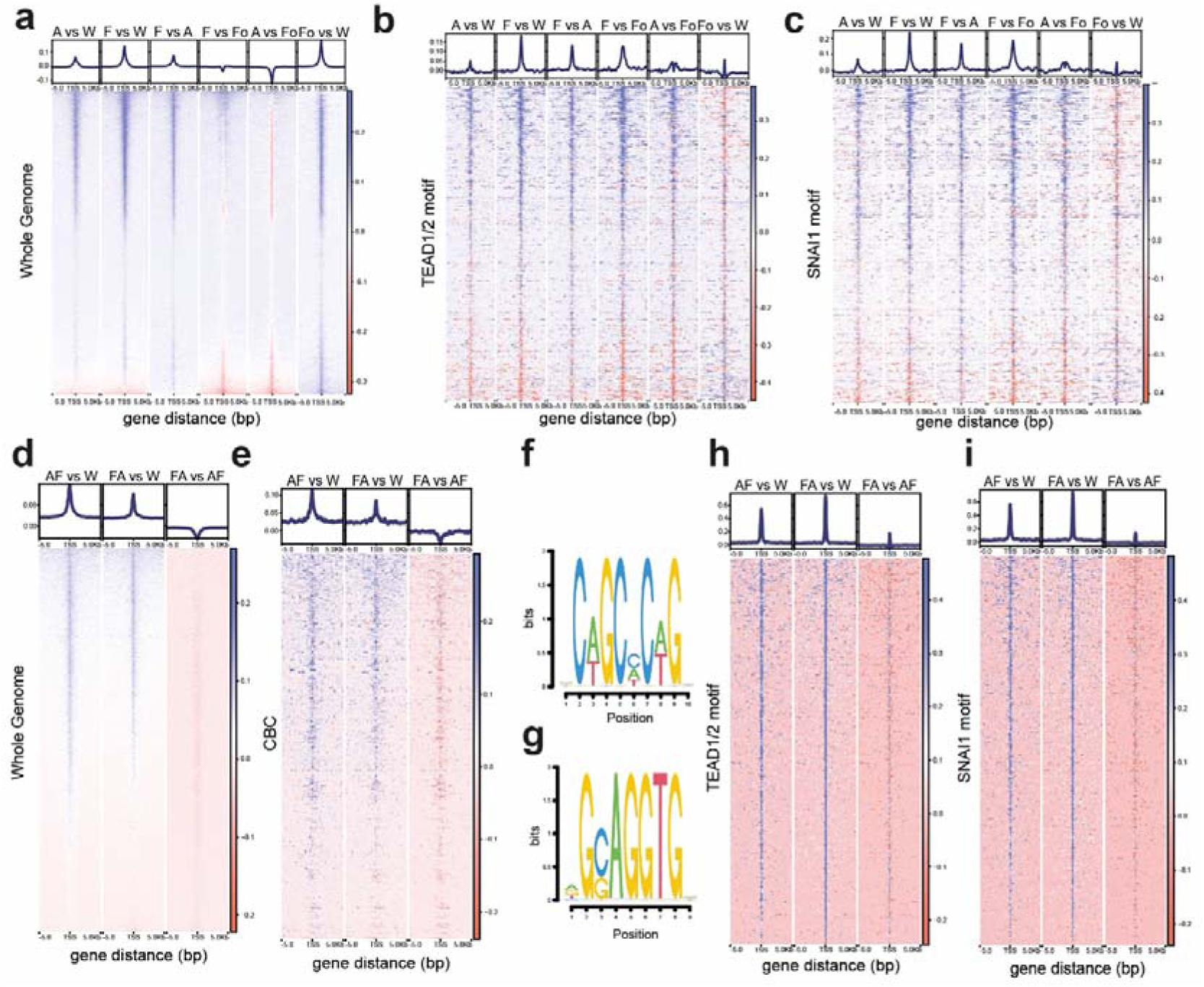
ATACseq analysis demonstrates cell slate changes are asscoiated with global and specific alterations in chromatin accessibility. **a,** Whole genome chromatin accessibility profiles for W, A, F and Fo organoids (blue = increased accessability, red = less accessability) **b,** Chromatin accessibility profiles *of* TEAD 1/2 motifs for W, A, F and Fo organoids (blue = increased accessability, red = less accessability). **c,** Chromatin accessibility profiles of SNAl1 motifs for W, A, F and Fo organoids (blue= increased accessability, red = less accessabillty). **d,** Whole genome chromatin accessibility profiles for F, AF and FA organoids (blue = increased accessability, red = less accessability). **e,** Chromatin accessibility profiles for CBC signature genes for W, AF and FA organoids (blue = increased accessability, red = less accessability). Motif plots of the two de novo motifs found enriched in the FA vs W organoid com­ parison were **(f)** “nCwGCmCwGn” and **(g)** “rGCAGGTGn”. Chromatin accessibility profiles of TEAD1/2 **(h)** and SNAl1 **(i)** motifs between W, AF and FA organoids. All experiments were performed with n=3 biological replicates with n=4 for foetal samples.

Next, we compared W, FA and AF organoids (matched for donor). In total, 14 543 peaks differed between AF and W organoids, 16 990 peaks differed between FA and W organoids, and 2 156 peaks differed between AF and FA organoids. Reassuringly, these findings suggest that there was much similarity in the effect of mutational order on chromosome accessibility between AF and FA. We noted that overall whole genome accessibility was reduced in FA vs W organoids compared with AF vs W organoids (Fig. 5d). Concordant with scRNAseq analysis between F and W organoids, we observed that there was increased accessibility to genes in the crypt base columnar (CBC) signature in AF vs W organoids compared with FA vs W organoids (Fig. 5e). De novo motif identification was performed as previous, and two motifs found enriched in the FA vs W organoid comparison included “nCwGCmCwGn” (Fig. 5f) and “rGCAGGTGn” (Fig. 5g). The top 3 closest matches to each of these motifs as per the JASPAR database, were *ZNF449*, *TEAD2*, and *TEAD1, SNAI1*, *SCRT2*, and *CTCFL* respectively. Intriguingly, TEAD1 and TEAD2 motifs were also more accessible in FA vs W organoids relative to AF vs W organoids (Fig. 5h). Finally, we observed increased chromatin accessibility of FA vs W organoids compared with AF vs W organoids at the SNAI1 transcription factor motif sites (Fig. 5i), which mediates EMT by repressing E-cadherin expression, therefore concurring with results observed in bulk-RNAseq and scRNAseq analysis of differentially expressed pathways. Taken together, our findings strongly implicate an epigenetic mechanism in which an early *FBXW7* mutation results in transcriptional repression of the consequences of a later *APC* mutation.

### Effects of mutation order are recapitulated in patient data

To perform human *in vivo* validation of our findings, we turned towards two datasets to infer the transcriptional impact of mutation order. First, we analysed transcriptional data from the recently published EPICC dataset [9]. Here, the authors performed matched single-crypt RNAseq and whole-genome sequencing on a small number of phenotypically normal crypts. We reanalysed transcriptional and genomic data from normal glands and performed PCA (Fig. 6a). Notably, one gland had an FBXW7-R578 mutation in the absence of an APC mutation, while another gland had an APC-S1346 mutation in the absence of an FBXW7 mutation. Two glands from the same patient possessed an identical FBXW7-R578 mutation as well as a separate APC-G1288 mutation; which can only be explained by the *FBXW7* mutation arising first, clonally expanding and then acquiring an *APC* mutation subsequently. Intriguingly, the gland with an *APC* mutation alone was located on PCA analysis distinct from all other glands. Notably, the two glands possessing both *APC* and *FBXW7* mutations also clustered closer to the wildtype glands and the single *FBXW7* mutant gland, analogous to our findings in gene-edited organoids. Whilst these inferences have been drawn based on a limited number of glands it is evident that FA mutations can also be found in phenotypically normal colon crypts.

**Fig. 6.**
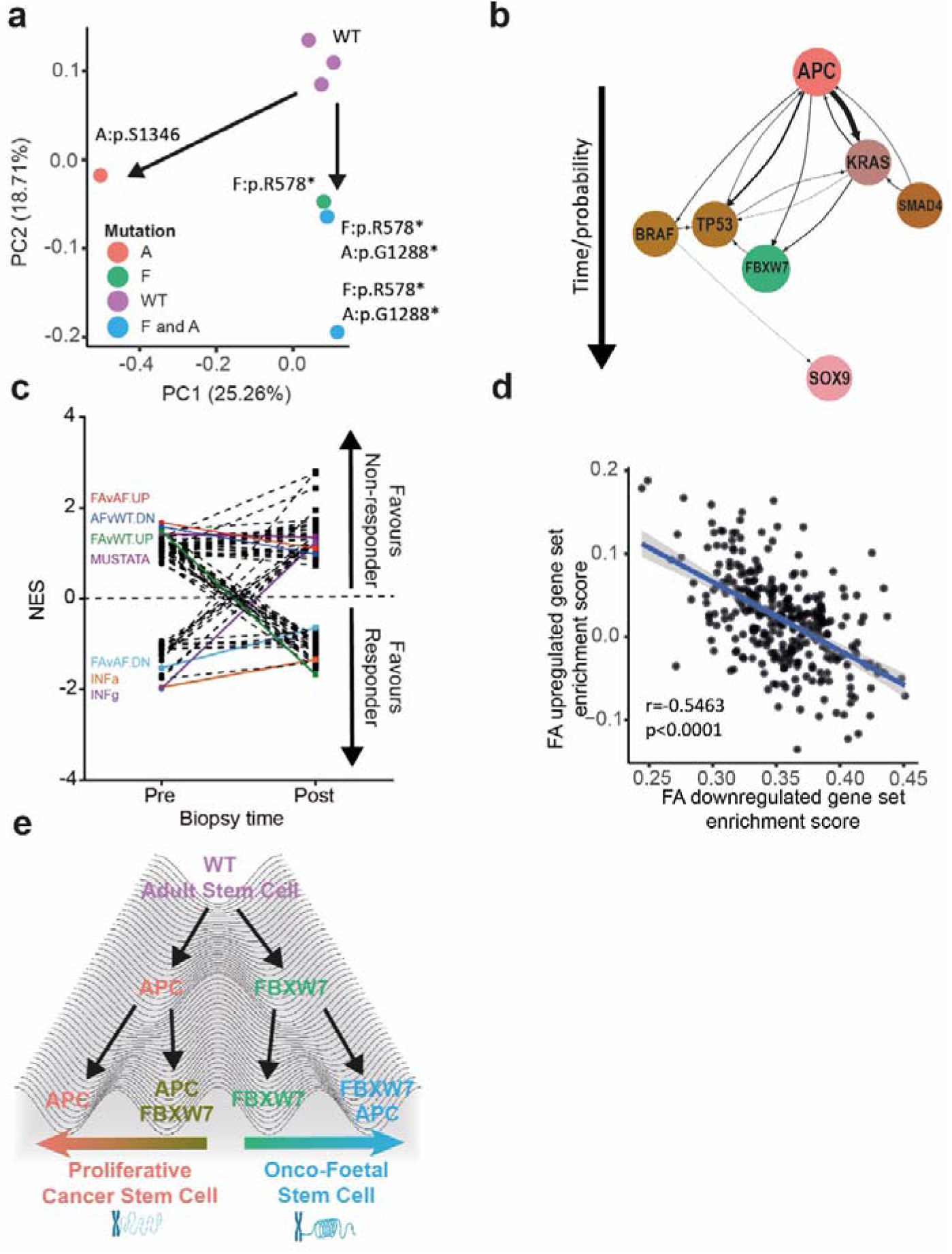
Patient data confirms FBXW7 mutation order-specific effects and identifies a novel chemotherapy predictive onco-foetal signature. **a,** PCA plot based on transcriptional grouping of ‘normal’ crypts containing F, A and no mutations. Mutations highlighted on plot. **b,** Plot representing ASCETIC analysis of 65 colonic adenomas from the S:CORT study. Arrow thickness represents strength of likelihood of mutation order for seven cancer genes sequenced. **c,** Linked column graph of GSEA NES for 73 Hallmark and stem cell signatures on FOxTROT trial pre and post treatment biopsies. Highlighted signatures significant on pre-treatment biopsies (p<0.05). **d,** Dot plot of enrichment scores in TCGA-COAD samples (n=320) for FA_UP and FA_DOWN signatures showing an inverse relationship (correlation score: −0.55, p<2.2 × 10·^15^). e, Waddington schematic of the consequences of mutation order on stem cell state. Adult stem cells (ISC/CBC) are driven down two axes: a proliferative cancer stem cell (proCSC) following APC mutation or an onco-foetal route following FBXW7 mutation.

Next, we adapted a newly described bioinformatic approach to infer mutational order from targeted sequencing data and applied this to a dataset (S:CORT) of 91 pre-malignant human colonic adenomas that had undergone targeted sequencing and transcriptional profiling [22] (Fig. 6b). Modified ASCETIC analysis showed that the invariant mode of adenoma development is an initial *APC* mutation (p<0.05). *FBXW7* mutations were never found to occur before *APC* in this cohort. In summary, whilst FA normal crypts can be observed in patient samples, our analysis of adenomas found that AF is the dominant mode of adenoma formation.

The foetal stem cell state in advanced malignancy has previously been associated with diminished response to chemotherapy [23, 24]. Chemotherapy can create somatic mutations in cancer and normal cells [25]. Given our findings that induced foetal states appear resistant to effects of further oncogenic transformation we hypothesised that the FA defined onco-foetal signature may also identify tumours resistant to chemotherapy. We analysed gene expression data from the Fluoropyrimidine, Oxaliplatin & Targeted Receptor pre-Operative Therapy for colon cancer study (FOxTROT), where patients were randomised to receive neoadjuvant chemotherapy prior to surgery [26]. Here, tumours were biopsied before and after treatment to identify transcriptional biomarkers of response. We generated differential gene expression lists by segregating patients into responders and non-responders with pre (n=95) and post (n=80) treatment biopsies. Next, we performed GSEA using the MSigDB Hallmarks and foetal gene signatures (n=73) unexpectedly finding the FA_up signature outperformed all previously published foetal and Hallmarks signatures in identifying non-responders to neo-adjuvant chemotherapy on pre-treatment biopsies (Fig. 6c and Extended Data Table 5). The AF signature as well as interferon response signatures on the other hand were strongly associated with improved responsiveness on pre-treatment biposies. Finally, ssGSEA of TCGA COAD/READ samples showed an inverse relationship between FA_up and FA_down gene signatures (correlation score: −0.55, *p*< 2.2 × 10^-16^) (Fig. 6e) [27]. Thus, whilst FA order of mutations in pre-neoplasia is an evolutionary dead end, the FA-defined onco-foetal signature in advanced malignancy is highly predictive of chemotherapy resistance.

## Discussion

Recent work has uncovered that molecular background in intestinal tumours influences a complex three-way interplay between revival/regenerative stem cell (foetal/YAP), homeostatic adult stem cell (CBC/ISC) and cancer stem cell populations (proCSC) [19, 21]. Whether similar processes play a role in pre-neoplasia and how this changes temporally remain unexplored. Using *FBXW7* as an example, our analyses demonstrate that epigenetics underlie the context-specific cellular response to somatic driver mutations, progressing the recent notion that tumour genotype alone does not equate to phenotype [28].

In our study, we found introducing an *FBXW7^-/-^* mutation in normal colon organoids generated little transcriptomic or proteomic response and organoids remained broadly ‘wildtype’ although primed towards a foetal state. These findings are compatible with the observation *in vivo* where *FBXW7* mutations are found in histologically normal crypts but failed to explain how *FBXW7* mutations are drivers in established CRC. In answer to this we uncovered an epistatic effect of the mutation when acquired on increasingly complex mutational backgrounds.

Building on our demonstration of epistasis we tested whether reordering mutations generated similar results. To our surprise not only were FA mutants phenotypically distinct from AF mutants but transcriptionally they also remained closer to the wildtype profiles, despite having lost two major tumour suppressor genes. Mutational order effects have been described once before in *JAK2/TET2* mutant haematological malignancies [29]. In a highly analogous manner, the authors found that blood cancers with switched order of mutations had alterations in stem cell state and transcriptomes.

To understand the link between mutation order, epistasis and stem cell state we tested whether an epigenetic mechanism could underly our data. Chromatin accessibility analyses showed that whilst both FA and AF organoids possessed more permissive chromatin than wildtype, broadly FA chromatin was less accessible than AF accounting for the general repression in the transcriptional consequences of an *APC* mutation. However, we found FA organoids did have increased accessibility to TEAD/SNAI1/AP-1 motifs. These data explain the foetal/YAP active state seen with F and FA mutations. Two very recent studies have explored chromatin landscapes in murine foetal and adult intestinal organoids [15, 30]. In highly concordant findings, these studies show similar TEAD/AP-1/SNAI1 family accessibility in foetal organoids and most importantly find *Fbxw7* to be a top hit in a CRISPR screen for regulators of the adult/foetal intestinal transition. Ultimately, the mechanism by which *FBXW7* mutations effect changes in chromatin accessibility remains cryptic, though a similar observation has been observed previously [31]. Given FBXW7’s important role in the E3 ubiquitin ligase complex, we speculate that decreased substrate degradation of one or a number of FBXW7 targets could result in this change in chromatin accessibility, though further work to elucidate this mechanism will need to be performed.

Our study develops on recent work describing Waddington-like plasticity in intestinal stem cell states [21]. We propose that whilst the normal colon crypt adopts predominantly the adult homeostatic stem cell state, altering the order of mutations affects subsequent fate trajectory. *FBXW7* mutation first cause cells to acquire a foetal state whereas an *APC* mutation first induces cells to acquire an additional proliferative/cancer stem cell state (Fig. 6f). These states remain epigenetically fixed and subsequent mutations prevent further plasticity between stem cell states/valleys. Overall, our study explains the basis for why *FBXW7* mutations in normal colonic epithelia fail to induce malignant transformation and provide the rationale for testing the translational impact of mutational order across a wider mutational spectrum and different cancers.

## Methods

### Human material for organoid cultures

Ethics approval for the retrieval of human colon specimens was accorded by the University of Oxford – Translation Gastroenterology Unit (16/YH/0247). Foetal samples were acquired under the HBDR project 200462, REC: 18/LO/0822. Written consent was obtained from all patients/donors.

### Organoid culture

Foetal organoids were derived from two donors (17pcw, Male, Proximal Colon) and (21pcw, Male, Proximal Colon). Adult colonic tissue was obtained from patients who were undergoing surgery for colorectal cancer. A 1 x 1 cm piece of colon from a region at least 5cm distant from the tumour, and appearing phenotypically normal was resected. Organoids were derived based on a previously published protocol [10]. Organoid culture growth media consists of advanced DMEM/F12 (Gibco), and supplemented with penicillin/streptomycin (Gibco), 10mM HEPES buffer solution (Gibco), 2mM GlutaMAX (Gibco), 50% Wnt3a conditioned medium (produced from ATCC CRL-2647 cell line), 25% R-spondin conditioned medium (produced from Cultrex HA-R-Spondin1-Fc 293T cells), 1X B-27 plus supplement (Gibco), 10μM SB 202190 (Tocris), 0.5μM SB 431542 (Tocris), 1μM prostaglandin E2 (Tocris), 50ng/ml Noggin (Peprotech), 50ng/ml EGF (Peprotech), 10mM nicotinamide (Sigma-Aldrich), and 1.25mM N-acetylcysteine (Sigma-Aldrich). For selection of APC mutants, Wnt3a and R-spondin conditioned media were withdrawn from the growth media for at least 3 weeks. For selection of TP53 mutants, growth media was supplemented with 10μM of Nutlin-3 for at least 1 month (Sigma-Aldrich).

### CRISPR/Cas9 gene editing of human organoids

sgRNAs were designed using Synthego’s CRISPR Design Tool (Extended Data Table 2). For *FBXW7*, a multiguide approach utilising three different guide RNAs in equal concentration was used. This increased the targeting efficacy given the lack of a selecting agent which can be used to enrich for *FBXW7* mutants. The *AAVS1* site is a commonly used safe harbour locus which served as a control to ensure studied effects were not a function of CRISPR gene editing.

Our protocol for gene editing human organoids has been described previously [10]. An electroporation-based approach was used to maximise gene editing efficacy. Cells were first dissociated into a single-cell suspension using TrypLe (Gibco) by incubating in a 37°C water bath for 10min. triturating at regular intervals. Organoids were then pelleted with centrifugation at 400g for 6 min. Pelleted organoids were resuspended in advanced DMEM, triturated and pelleted by centrifugation once more. A ribonucleoprotein (RNP) complex comprising Cas9 (Sigma-Aldrich) and guide RNA in a molar ratio of 1:4 was resuspended in Buffer R (Invitrogen) and left to stand at room temperature for 10min. Pelleted organoids were resuspended in Buffer T (Invitrogen). The RNP complex and resuspended organoids were then mixed. Electroporation was performed using a Neon Transfection System (ThermoFisher) with the following settings – voltage 1350V, width 20ms, and pulses 2. After electroporation, organoids are transferred into advanced DMEM and left to recover for 10min. Centrifugation was performed to pellet organoids, which were then resuspended in BME and transferred to cell culture plates as described above. Following electroporation, organoids were incubated for one week and allowed to expand. Primers which generated amplicons of approximately 500bp in length were designed on Benchling (Extended Data Table 2). Amplicons were Sanger sequenced by an external vendor (Genewiz). Efficiency of targeting was carried out using Synthego Performance Analysis, ICE Analysis. 2019. V3.0 Synthego.

Human colon organoids routinely senesce ∼ P7/8 necessitating continual re-derivation, targeting and expansion. In all experiments each replicate represents a distinct clone generated from an individual patient unless stated otherwise. Each edited clone was used for assays once targeting efficiency through selection (APC/TP53) or repeated targeting (FBXW7) was >80%.

### Protein lysates and western blot

Proteins were harvested using RIPA (ThermoFisher) with addition of protease and phosphatase inhibitor cocktails (Sigma-Aldrich). Quantification of protein concentration was performed using BCA assay (ThermoFisher).

For western blot, 30µg of protein was used for each lane. Proteins were separated using a NuPage 4 – 12% Bis-Tris gel (Invitrogen) and transferred onto a 0.45µm nitrocellulose membrane. For phosphorylated proteins, 5% BSA was used. For non-phosphorylated proteins, 5% low-fat milk was used. The membrane was blocked for 1 hour before overnight incubation with primary antibodies at 4°C. Rabbit anti-human FBXW7 antibody (1:2500, BS-8394R, Bioss, USA), rabbit anti-human phosphor-CJUN antibody (1:2500, PA5-40193, Invitrogen, USA), rabbit anti-human CJUN antibody (1:2500, ab40766, Abcam, USA), rabbit anti-human phosphor-CCNE1 antibody (1:2500, ab52195, Abcam, USA), rabbit anti-human CCNE1 antibody (1:2500, ab33911, Abcam, USA), rabbit anti-human phosphor-CMYC (1:2500, ab185655 and ab185656, Abcam, USA), rabbit anti-human CMYC (1:2500, ab32072, Abcam, USA), rabbit anti-human NICD (1:2500, #4147, Cell signalling technology, USA), rabbit anti-human NOTCH1 (1:2500, ab52627, Abcam, USA), rabbit anti-human beta-catenin (1:1000, #8480, Cell signalling technology, USA), rabbit anti-human p53 (1:1000, #9282, Cell signalling technology, USA) and rabbit anti-human GAPDH (1:2500, ab52627, Abcam, USA) was used. The membrane was then incubated with goat anti-rabbit antibody (1:5000, ab6721, Abcam, USA) at room temperature for 1 hour. The membranes were then imaged with a ChemiDoc XRS+ system (Bio-Rad).

### Immunofluorescence and imaging

Immunofluorescence on organoids was previously described in our protocol [10]. Briefly, media from the wells of organoids was removed and fixation with 2% paraformaldehyde was performed. This was incubated at room temperature for 30min. Organoids were then washed with ice-cold DPBS gently and allowed to settle by gravity. The supernatant was then removed and blocked with Organoid Blocking Solution comprising DPBS supplemented with 3% BSA, 1% Triton X-100, 1% saponin and 1% secondary antibody animal serum. Blocking was performed for 3 hours at room temperature. Organoids were again allowed to settle by gravity, and the supernatant was removed. Primary antibody diluted in organoid blocking solution was then added to the organoids for 24 hours at 4°C. Organoids were then washed with DPBS supplemented with 3% BSA, 1% Triton X-100, and 1% saponin. Secondary antibody diluted in the organoid washing solution was then added to the organoids for 2 hours at 4°C. After 2 hours, organoids were again washed with organoid washing solution. Phalloidin stain was added to the organoids for 60min, before DAPI stain was added to the organoids for 5min. Imaging was performed with the Andor Dragonfly High-Speed Confocal Microscope Systems (Oxford Instruments). Antibodies used were mouse anti-human FBXW7 antibody (1:100, H00055294-M02. Novus Biologicals, USA), rabbit anti-LY6D goat anti-mouse (1:200, 17361-1-AP, Proteintech, USA), IgG H&L (Dylight 550) (1:500, ab96872, Abcam, USA), IgG H&L (Alexa Fluor 488) (1:1000, Abcam, USA), Phalloidin iFluor 647 (1:400, ab176759, Abcam, USA), and DAPI (300nM, D1306, Invitrogen, USA).

### Co-culture of F, A and W organoids

Transduction of organoids was performed to achieve stable expression of GFP or mCherry fluorescence. Lentiviral particles were first generated by transfection of HEK293T cells with pMDLg/pRRE, pRSV-Rev, pSF-CMV-VSVG (Sigma-Aldrich) and either pCDH-EF1-copGFP-T2A-Puro for GFP fluorescence or pCDH-EF1-copGFP-T2A-Puro for mCherry fluorescence. pMDLg/pRRE, and pRSV-Rev were gifts from Didier Trono (Addgene plasmid # 12251; http://n2t.net/addgene:12251; RRID:Addgene_12251) [32], pCDH-EF1-copGFP-T2A-Puro and pCDH-CMV-mCherry-T2A-Puro were gifts from Kazuhiro Oka (Addgene plasmid # 72263; http://n2t.net/addgene:72263; RRID:Addgene_72263).

To transduce organoids with the lentivirus particle, the organoids were first dissociated into single cells by TrypLe for 10min in a 37°C water bath, with trituration at regular intervals. Single cells were centrifuged at 400g for 6 min to pellet cells, which were then passed through a 50μm filter. The number of cells was then counted using a hemocytometer, and the required number of cells was aspirated, centrifuged and then pelleted. These cells were then resuspended in organoid growth media, and viral particles were added to achieve a multiplicity of infection (MOI) of 20. Cells were infiltrated over wells which had been overlaid with a pre-polymerised BME layer and left in a humidified 37°C incubator for 24 hours. After 24 hours, dead cells were found floating above the BME, while live cells would have become embedded in the BME. The media and dead cells were aspirated and fresh organoid growth media was added. Fluorescence was checked using an EVOS M5000 cell imaging system (ThermoFisher) after 72 hours.

For the co-culture of organoids, a GFP-labelled population was co-cultured with an mCherry-labelled population in equal amounts to ascertain differences in proliferation. Cell culture wells were imaged on day 0, 1, 3, 5 and 7 using an EVOS M5000 cell imaging system (ThermoFisher). ImageJ was used to determine fluorescent intensity on each of these days.

### Bulk RNA sequencing and analysis

Extraction of total RNA was performed using the RNeasy Mini Kit (Qiagen) as per manufacturer’s protocol. The quantity and quality of the RNA was ascertained using the Agilent 2100 Bioanalyzer. Sequencing was performed by the Oxford Genomics Centre, Oxford using an Illumina NextSeq500 instrument using the standard paired-end protocol with a read length of 150bp. Fastq reads were processed to clip low quality leading and trailing edges and to remove any adapter content using Cutadapt (v.3.5). Quality checked FASTQ reads were aligned to the GRCh38 human genome and gene annotation (Ensembl release 105) using STAR aligner (v.2.7.3a) two-pass mode to generate gene-level quantification. Raw counts were processed in R (v.4.1.3) for all statistical testing and plotting purposes. Normalisation and differential expression were performed using limma (v.3.50.3). This included the generation of heatmaps using gplots (v.3.1.1), volcano plots using ggplot2 (v3.3.5) and PCA plots using ggfortify (v.0.4.14). Gene set enrichment analysis was performed using fgsea (v.1.20.0) and GSVA (v.1.42.0).

### Single-cell RNA sequencing and analysis

scRNA-seq was conducted using a 5’ scRNA-seq gene expression workflow (10x Genomics). Cells were loaded onto a 10x Chromium Controller for GEM generation followed by single cell library construction using 10x Chromium Next GEM Single Cell 5’ Library and gel bead kit v1.1 following manufacturer’s instructions. Size profiles of amplified cDNA and final libraries for sequencing were verified by electrophoresis (Agilent 2100 Bioanalyzer system, High Sensitivity DNA Kit), and the concentration of final libraries was measured with Qubit (Thermo Fisher Scientific). Libraries were sequenced on an Illumina NextSeq 2000 (26 cycles read 1, 8 cycles i7 index, 98 cycles read 2), achieving a mean of 39,412 reads per cell. Gene by cell barcode quantification matrix were obtained using Cell Ranger (v.7.0.1) and Seurat (v.4.2.0) was used to perform normalisation and UMAP based cluster assignment of pass cells.

Single sample gene set enrichment analyses (ssgsea) method from GSVA (v.1.42.0) was used on a custom list of gene signature sets defined by editing the MSigDb (v.7.5.1) pathways to leave out SPERMATOGENSIS, MYOGENESIS and PANCREAS_BETA_CELLS, and replacing the default WNT_BETA_CATENIN_SIGNALING set by WNT_SIGNALING gene set (MSigDb systematic name M5493), as per Househam et al. [33]. Additional gene sets included were – genes up- or down-regulated in bulk RNA-seq comparison of ‘FBXW7-APC’ versus ‘APC-FBXW7’ organoids (adjusted p-value < 0.05 and logFC >= |1.5|; n= 248 up- and n= 166 down-regulated genes), mouse foetal gastric epithelium gene set from Vallone et al. 2016 (n= 122 human homologues), intestinal stem cell signature from Merlos-Suárez et al. 2011 (n= 74), YAP pathway signature from Gregorieff et al. 2015 (n= 213 human homologues), crypt-base columnar (CBC) and regenerative stem cell (RSC) signature sets from Vazquez et al. 2022 (n= 340 and 206 human homologues respectively)(Extended Data Table 3). Enrichment scores obtained per gene set was z-score transformed, and cells with scores greater than 3^rd^ quartile or less than 1^st^ quartile of the score distribution were termed as enriched or depleted respectively, for that gene set. To assess statistical enrichment or depletion of a given gene set in FBXW7 mutant versus wildtype categories, chi-square test was applied on contingency table for cell counts from enriched or depleted classes. Pearson residuals were visualised using mosaic plot from R package vcd (v.1.4-11). For RNA velocity analysis, separate count matrices for spliced and unspliced transcripts were created using Kallisto-Bustools (v.0.27.3) with the La Manno et al. 2018 strategy, while using the same reference annotation from Cell Ranger (GRCh38, v.2020-A). Spliced/ unspliced count data were combined with Seurat based UMAP clustering using scVelo (v.0.2.5) under python v.3.10.6. RNA velocity graph computed was overlaid on UMAP cluster embeddings to infer trajectory direction.

### Proteomics

Four replicates per condition, each containing 50 µg protein, were solubilised in 5% SDS then processed by S-Trap micro (Protifi) protocol according to the manufacturer’s instructions. Digestion was performed overnight with a 1:25 ratio of trypsin (Sequencing Grade, Promega) to protein. Tryptic peptides were dried by vacuum centrifugation, then reconstituted prior to MS analysis in 0.1% formic acid. LC-MS/MS analysis was performed using the Orbitrap Ascend Tribrid instrument (Thermo Scientific) connected to a Thermo Scientific Vanquish^TM^ Neo UHPLC system interfaced using a nano-EASY spray source. The Vanquish^TM^ Neo was operated in Trap and Elute mode using 0.1 % Formic acid in water as solvent A, 0.1% formic acid in acetonitrile as solvent B and strong wash buffer and 0.1% trifluoroacetic acid as weak wash. Tryptic peptides were loaded onto a PepMap Neo C18 Trap (Thermo Scientific; S/N 174500) at 8 ul/min (total volume of 24ul) and separated on a 50C heated EasySpray Pep Map Neo (Thermo Scientific; S/N ES75500) column using a multistep gradient going from 2% to 18% solvent B in 40 min and from 18% to 35% solvent B in 20 min at 300nl/min flow rate. The column was washed for 14 min with 99% B, followed by the fast equilibration on the Vanquish Neo (combine control mode, upper pressure to 1000bar). In parallel, the trap was subjected to the zebra wash and fast equilibrated.

MS data were acquired in data-independent mode (DIA) with minor changes from previously described method [34–36]. Briefly, MS1 scans were collected in the orbitrap at a resolving power of 45K at m/z 200 over m/z range of 350 – 1650m/z. The MS1 normalised AGC was set at 125% (5e5ions) with a maximum injection time of 91 ms and a RF lens at 30%. DIA MS2 scans were then acquired using the tMSn scan function at 30K orbitrap resolution over 40 scan windows with variable width, with a normalized AGC target of 1000%, maximum injection time set to auto and a 30 % collision energy.

Raw data were searched in DIA-NN v1.8.1 in library-free mode against the UniProt human proteome database (UP000005640, downloaded 18^th^ May 2023), plus common contaminants [37]. A maximum of one missed cleavage was permitted for Trypsin/P digestion, with cysteine carbamidomethylation set as a fixed modification. ‘Match between runs’ and retention time-dependent cross-run normalisation were enabled, with mass accuracy settings inferred from the data. The DIA-NN neural network classifier was set to double-pass mode. Further data analysis was performed in Perseus v2.0.11, where protein groups were filtered to include only those identified in at least three replicates of one experimental condition. Protein intensity values were log2-transformed, then missing values were imputed with random values generated from a downshifted normal distribution (width 0.3, downshift 1.8). Data are available via ProteomeXchange with identifier PXD052352.

To identify differential protein expression Student’s t-test was applied on replicate level data from *FBXW7* mutant and WT samples, for each of the detected proteins using the “t.test” function. The p-values thus obtained were corrected for multiple-testing using the Benjamini and Hochberg (i.e. FDR) method by using the “p.adjust” function. All calculations were done in R ver. 4.3.3.

### Assay for transposase-accessible chromatin sequencing (ATACseq) and analysis

Human colon organoids were generated as previously described and suspended in 40ul of BME in 400ul of complete organoid media. At Passage 2, organoids were subjected to electroporation based CRISPR using either no guide, multiguide RNA targeting FBXW7 or single guide RNA targeting APC (ref STAR protocols). Organoids targeted with guide RNA against APC were subsequently grown in complete organoid media (also in STAR paper) omitting WNT3a conditioned media until 7 days prior to ATAC-sequencing. Organoids targeted with FBXW7 guide RNA underwent a second round of electroporation based CRISPR at passage 3 to achieve >80% knockout of FBXW7. Wildtype and knockout organoids were subsequently passaged to achieve 1 × 10^12^ cells, grown in identical complete organoid media for 7 days and submitted for library preparation and ATAC-sequencing (Genewiz/Azenta). ATAC experiments were carried out separately for W, A, F and Foetal (1) and W, AF and FA (2). To ensure that background epigenetic status was corrected for, all CRISPR-edited organoids for these experiments were generated from the same donor W organoids for each of the two experiments.

Library preparation for W/FA/AF ATACseq was performed using the ATAC-Seq Kit (Active Motif) as per the manufacturer’s instructions. Multiplexing was performed using unique i7 and i5 indexed primers. The quantity and quality of DNA was assessed using the Agilent High Sensitivity DNA Kit, ensuring that transposed DNA fragments were between 200 and 1000bp, with a periodicity of ∼200bp. Sequencing was performed by the Oxford Genomics Centre, Oxford using an Illumina NextSeq2000 instrument.

Quality checked fastq reads were aligned to the human genome (GRCh38) using bwa aligner (v.0.7.17). Alignment output (BAM files) were filtered, and TA (Tag Alignment) format data created using samtools (v.1.14), picard (v.2.6.0) and bedtools (v.2.30.0). Further, MACS3 (v.3.0.0b2) was used to perform peak calling which were then annotated using ChIPseeker (v.1.30.3). MACS3 identified peaks for the ‘AF vs W’ and ‘FA vs W’ comparisons were thresholded to select those with score (i.e., −10*log(q-value)) >median of the respective distribution, overlapping regions merged and the 50bp span from the centre of the peak region were extracted as input for *de novo* transcription factor motif enrichment analysis using rGADEM (v.2.46.0). Identified motifs were plotted using TFBSTools (v.1.36.0) and annotated for the closest matching core human transcription factors using JASPAR2020 (v.0.99.0). Gene/ motif-set level visualisation of ATAC-seq signal enrichment was performed using deeptools (v.3.5.2). Comparative bigwig files were created for ‘AF vs W’, ‘FA vs W’ and ‘FA vs AF’ with BPM (Bins per million) normalisation, non-covered regions skipped, ENCODE blacklist regions filtered out and effective genome size adjusted to reflect the filtering. The bigwig files were then profiled for ATAC-seq signal enrichment with reference-point at the TSS (Transcription Start Site) and upstream/ downstream 5kb region. Signal enrichments were profiled for each of these sets –all genes in the genome (gencode v.38; ‘Whole Genome’), the CBC gene signature set (‘CBC’) and the 50bp genomic regions where de novo motif search identified TEAD1/2 motifs (542 regions; ‘TEAD1/2 motif’) or SNAI1 motifs (491 regions; ‘SNAI1 motif’).

Comparative bigwig files were created from the final BAM files for ‘F’, ‘A’ and ‘Foetal’ organoids by comparing each to the ‘WT’ organoid BAM, using ‘bamCompare’ from deeptools (v.3.5.2) with parameters: binSize 10, normalisation BPM, ENCODE blacklist regions filtered out and effective genome size adjusted to reflect the filtering. The bigwig files were then used to create the accessibility tracks by extracting UCSC Genome Browser data (ver. GRCh38) and plotting it in R (v.4.3.3) using bwtool (v.1.0) and trackplot (v1.5.10).

### Validation of results in published patient datasets

Three independent published datasets were used to validate our findings. Somatic mutation and gene expression for the TCGA-COAD cohort were obtained from NCI-GDC data portal [27], to create a dataset of colon adenocarcinoma cases only from 320 unique patients. Per-sample enrichment scores for the gene sets was calculated using ‘ssgsea’ method in GSVA (v.1.42.0). The second validation made use of transcriptomic results derived from the EPICC dataset [9]. Here, individual gland-level transcriptomic results were obtained. Generation of PCA plots using ggplot2 (v3.3.5). The third S:CORT study provided clinical and molecular data for both ASCETIC analysis of adenomas and transcriptional analysis of predictors of neo-adjuvant chemotherapy (FOxTROT) [26].

### Modified ASCETIC analysis

The Agony-based cancer evolution inference (ASCETIC) framework was employed to identify conserved patterns of driver gene mutations across a cohort of bulk sequenced adenomas [22]. For 91 adenomas, copy number and ploidy estimate were used to estimate cancer cell fraction of each single nucleotide variant at the single sample level. Functions from the ASCETIC package were used to calculate a partial ordered set of genes based on minimising agony across the entire cohort and generate a ranking estimate of genes containing single nucleotide variants. Genes containing the most commonly occurring single nucleotide variants were displayed graphically, with the y-axis representing overall probability of early mutation, based on recurrent high confidence (p<0.05) evolutionary steps and arrow thickness, representing recurrent evolutionary step number (Fig. 6b).

## Acknowledgements

S.J.A.B. was supported by The Pharsalia Trust, UK & a Cancer Research UK Advanced Clinician Scientist Fellowship (C14094/A27178). D.K.H.C. was supported by Singapore Ministry of Health’s National Medical Research Council under a research training fellowship (MOH-FLWSHP10may-0001). A.S. was supported by the UK Medical Research Council, Wellcome Investigator Award (219523/Z/19/Z). D.F-C was supported by an NIHR Academic Clinical Lectureship Award. The Stratification in Colorectal Cancer Consortium (S:CORT) was funded by the Medical Research Council and Cancer Research UK (MR/M016587/1). The human foetal material was provided by the Joint MRC/Wellcome Trust (MR/R006237/1) Human Developmental Biology Resource (www.hdbr.org) and we thank B. Crespo, Professor Copp and the entire HDBR team.

## Author contributions

D.K.H.C. and S.J.A.B. conceived the overall experimental questions and design. D.K.H.C. performed experiments, wrote the initial draft of the manuscript, and was supervised by S.J.A.B. A.M. and S.J.A.B. performed bioinformatic analysis for the project. S.D.C. assisted with immunofluorescence of organoids. Y. Z. generated plasmids for lentiviral labelling of organoids, performed the proteomics experiment and W vs APC scRNAseq experiment. R.O. H.F. S.J. and X.L. assisted with scRNAseq. J.H. and T.G. contributed the EPICC dataset and assisted with its analysis. J.B. performed the ASCETIC and EPICC analysis and carried out the W, F and A ATACseq experiment. I.V. S.F. and R.F. carried out proteomic analysis. D.F.C and A.S. assisted with foetal organoid experiments. S.J.A.B. was overall in-charge of the project. All authors contributed to drafting and final approval of the manuscript.

## Materials and correspondence

Raw sequence data would be made available via the European Genome-Phenome Archive (EGA) and proteomics via PRIDE, on publication of the manuscript. The mass spectrometry proteomics data have been deposited to the ProteomeXchange Consortium via the PRIDE partner repository with the dataset identifier PXD052352. Please relay correspondence to Simon Buczacki.

## Inclusion and ethics in global research

Research was conducted in accordance with principles of inclusion and good ethics in global research.

## Code availability

All relevant scripts to replicate the analysis would be made available on Github on publication of the manuscript.

