## Extended data tables for "Mutational order and epistasis determine the consequences of *FBXW7* mutations during colorectal cancer evolution"

### Extended Data Table 1 Reactome pathway analysis of F vs W proteomics

| pathway | pval | padj | log2err | ES | NES | size |
|---|---|---|---|---|---|---|
| REACTOME_RHO_GTPASES_ACTIVATE_PKNS | 1.98E-08 | 1.61E-06 | 0.734 | -0.811 | -2.396 | 34 |
| REACTOME_POSITIVE_EPIGENETIC_REGULATION_OF_RRNA_EXPRESSION | 2.37E-08 | 1.61E-06 | 0.734 | -0.750 | -2.371 | 47 |
| REACTOME_B_WICH_COMPLEX_POSITIVELY_REGULATES_RRNA_EXPRESSION | 4.49E-09 | 6.1E-07 | 0.761 | -0.811 | -2.359 | 33 |
| REACTOME_ERCC6_CSB_AND_EHMT2_G9A_POSITIVELY_REGULATE_RRNA_EXPRESSION | 8.06E-09 | 9.13E-07 | 0.748 | -0.850 | -2.328 | 25 |
| REACTOME_ACTIVATION_OF_ANTERIOR_HOX_GENES_IN_HINDBRAIN_DEVELOPMENT_DURING_EARLY_EMBRYOGENESIS | 1.11E-07 | 5.4E-06 | 0.705 | -0.777 | -2.309 | 35 |
| REACTOME_FORMATION_OF_THE_BETA_CATENIN_TCF_TRANSACTIVATING_COMPLEX | 5.81E-08 | 3.04E-06 | 0.720 | -0.832 | -2.278 | 25 |
| REACTOME_MEIOTIC_RECOMBINATION | 2.36E-08 | 1.61E-06 | 0.734 | -0.869 | -2.262 | 20 |
| REACTOME_INHIBITION_OF_DNA_RECOMBINATION_AT_TELOMERE | 4.04E-07 | 1.72E-05 | 0.675 | -0.835 | -2.225 | 22 |
| REACTOME_PRC2_METHYLATES_HISTONES_AND_DNA | 4.98E-08 | 2.82E-06 | 0.720 | -0.903 | -2.222 | 15 |
| REACTOME_RUNX1_REGULATES_GENES_INVOLVED_IN_MEGAKARYOCYTE_DIFFERENTIATION_AND_PLATELET_FUNCTION | 2.06E-06 | 6.36E-05 | 0.627 | -0.817 | -2.219 | 23 |
| REACTOME_BASE_EXCISION_REPAIR | 1.18E-06 | 4.01E-05 | 0.644 | -0.732 | -2.213 | 38 |
| REACTOME_RNA_POLYMERASE_I_PROMOTER_ESCAPE | 1.09E-06 | 4.01E-05 | 0.644 | -0.758 | -2.205 | 33 |
| REACTOME_NEGATIVE_EPIGENETIC_REGULATION_OF_RRNA_EXPRESSION | 4.47E-06 | 0.000119 | 0.611 | -0.699 | -2.193 | 44 |
| REACTOME_RNA_POLYMERASE_I_TRANSCRIPTION | 5.13E-06 | 0.00012 | 0.611 | -0.686 | -2.187 | 49 |
| REACTOME_DEPOSITION_OF_NEW_CENPA_CONTAINING_NUCLEOSOMES_AT_THE_CENTROMERE | 7.49E-07 | 2.99E-05 | 0.659 | -0.867 | -2.166 | 17 |
| REACTOME_TELOMERE_MAINTENANCE | 5.09E-06 | 0.00012 | 0.611 | -0.678 | -2.165 | 50 |
| REACTOME_CONDENSATION_OF_PROPHASE_CHROMOSOMES | 2.32E-06 | 6.87E-05 | 0.627 | -0.850 | -2.157 | 18 |
| REACTOME_ASSEMBLY_OF_THE_ORC_COMPLEX_AT_THE_ORIGIN_OF_REPLICATION | 3.24E-06 | 9.18E-05 | 0.627 | -0.846 | -2.146 | 18 |
| REACTOME_BASE_EXCISION_REPAIR_AP_SITE_FORMATION | 1.78E-06 | 5.78E-05 | 0.644 | -0.872 | -2.145 | 15 |
| REACTOME_CHROMOSOME_MAINTENANCE | 4.59E-06 | 0.000119 | 0.611 | -0.657 | -2.134 | 56 |
| REACTOME_RUNX1_REGULATES_TRANSCRIPTION_OF_GENES_INVOLVED_IN_DIFFERENTIATION_OF_HSCS | 5.3E-06 | 0.00012 | 0.611 | -0.650 | -2.132 | 61 |
| REACTOME_TRANSCRIPTIONAL_REGULATION_BY_SMALL_RNAS | 5.82E-06 | 0.000128 | 0.611 | -0.663 | -2.129 | 52 |
| REACTOME_PHASE_II_CONJUGATION_OF_COMPOUNDS | 1.07E-05 | 0.00022 | 0.593 | -0.665 | -2.126 | 51 |
| REACTOME_HDACS_DEACETYLATE_HISTONES | 4.72E-06 | 0.000119 | 0.611 | -0.737 | -2.120 | 31 |
| REACTOME_NONHOMOLOGOUS_END_JOINING_NHEJ | 1.72E-05 | 0.000325 | 0.576 | -0.742 | -2.111 | 29 |
| REACTOME_DISEASES_OF_PROGRAMMED_CELL_DEATH | 1.9E-05 | 0.000348 | 0.576 | -0.679 | -2.099 | 42 |
| REACTOME_NON_INTEGRIN_MEMBRANE_ECM_INTERACTIONS | 6.36E-05 | 0.000939 | 0.538 | -0.710 | -2.099 | 34 |
| REACTOME_MEIOSIS | 5.35E-05 | 0.000809 | 0.557 | -0.700 | -2.093 | 36 |
| REACTOME_OXIDATIVE_STRESS_INDUCED_SENESCENCE | 2.3E-05 | 0.000412 | 0.576 | -0.671 | -2.086 | 43 |
| REACTOME_LAMININ_INTERACTIONS | 5.21E-05 | 0.000806 | 0.557 | -0.768 | -2.086 | 23 |
| REACTOME_PRE_NOTCH_EXPRESSION_AND_PROCESSING | 3.44E-05 | 0.000571 | 0.557 | -0.699 | -2.075 | 35 |
| REACTOME_RHO_GTPASE_EFFECTORS | 3.85E-08 | 2.38E-06 | 0.720 | -0.535 | -2.040 | 166 |
| REACTOME_EPIGENETIC_REGULATION_OF_GENE_EXPRESSION | 9.08E-06 | 0.000193 | 0.593 | -0.588 | -2.035 | 80 |
| REACTOME_TRANSCRIPTIONAL_REGULATION_OF_GRANULOPOIESIS | 8.44E-05 | 0.001222 | 0.538 | -0.760 | -2.025 | 22 |
| REACTOME_RRNA_MODIFICATION_IN_THE_NUCLEUS_AND_CYTOSOL | 3.29E-05 | 0.000568 | 0.557 | -0.617 | -2.016 | 58 |
| REACTOME_AMYLOID_FIBER_FORMATION | 0.000154 | 0.001766 | 0.519 | -0.655 | -2.003 | 40 |
| REACTOME_MEIOTIC_SYNAPSIS | 9.83E-05 | 0.001364 | 0.538 | -0.729 | -2.000 | 26 |
| REACTOME_THE_CITRIC_ACID_TCA_CYCLE_AND_RESPIRATORY_ELECTRON_TRANSPORT | 3.94E-07 | 1.72E-05 | 0.675 | 0.516 | 2.054 | 138 |
| REACTOME_MITOCHONDRIAL_TRANSLATION | 1.14E-06 | 4.01E-05 | 0.644 | 0.562 | 2.087 | 90 |
| REACTOME_RESPIRATORY_ELECTRON_TRANSPORT | 2.77E-09 | 4.71E-07 | 0.775 | 0.648 | 2.387 | 85 |
| REACTOME_RESPIRATORY_ELECTRON_TRANSPORT_ATP_SYNTHESIS_BY_CHEMIOSMOTIC_COUPLING_AND_HEAT_PRODUCTION_BY_UNCOUPLING_PROTEINS | 5.77E-10 | 1.96E-07 | 0.801 | 0.628 | 2.391 | 98 |
| REACTOME_COMPLEX_I_BIOGENESIS | 2.26E-09 | 4.71E-07 | 0.775 | 0.735 | 2.395 | 47 |

**Extended Data Table 2 Primer and guide sequences for gene editing**

| Gene target | guide RNA sequences | Forward primer | Reverse primer |
|---|---|---|---|
| *FBXW7* | GCAAGGAATGGTGAAGTTGT<br>GATGAATCGTGTGGTAGAGG<br>AGCAAAAGACGACGAACTGG | TTTTCCCCTGCAGAATGTGA | ACTGGAGTTCGTGACACTGTT |
| *APC* | UCUGUAUAAAUGGCUCAUCG | ATGCTGCAGTTCAGAGGGTC | TTTTTCTGCCTCTTTCTCTTGGT |
| *TP53* | CCGGUUCAUGCCGCCCAUGC | CTTGCCACAGGTCTCCCCAAG | AGCCACAGGTTAAGAGGTCC |
| AAVS1(*PPP1R12C*) | GGGGCCACUAGGGACAGGAU | GGTCCGAGAGCTCAGCTAGT | GGCTCCATCGTAAGCAAACC |

### Extended Data Table 3 Gene signatures used for GSEA and ssGSEA

| Pathway/ Gene-set name | Source |
|---|---|
| TNFA_SIGNALING_VIA_NFKB | MSigDb.Hallmark |
| HYPOXIA | MSigDb.Hallmark |
| CHOLESTEROL_HOMEOSTASIS | MSigDb.Hallmark |
| MITOTIC_SPINDLE | MSigDb.Hallmark |
| TGF_BETA_SIGNALING | MSigDb.Hallmark |
| IL6_JAK_STAT3_SIGNALING | MSigDb.Hallmark |
| DNA_REPAIR | MSigDb.Hallmark |
| G2M_CHECKPOINT | MSigDb.Hallmark |
| APOPTOSIS | MSigDb.Hallmark |
| NOTCH_SIGNALING | MSigDb.Hallmark |
| ADIPOGENESIS | MSigDb.Hallmark |
| ESTROGEN_RESPONSE_EARLY | MSigDb.Hallmark |
| ESTROGEN_RESPONSE_LATE | MSigDb.Hallmark |
| ANDROGEN_RESPONSE | MSigDb.Hallmark |
| PROTEIN_SECRETION | MSigDb.Hallmark |
| INTERFERON_ALPHA_RESPONSE | MSigDb.Hallmark |
| INTERFERON_GAMMA_RESPONSE | MSigDb.Hallmark |
| APICAL_JUNCTION | MSigDb.Hallmark |
| APICAL_SURFACE | MSigDb.Hallmark |
| HEDGEHOG_SIGNALING | MSigDb.Hallmark |
| COMPLEMENT | MSigDb.Hallmark |
| UNFOLDED_PROTEIN_RESPONSE | MSigDb.Hallmark |
| PI3K_AKT_MTOR_SIGNALING | MSigDb.Hallmark |
| MTORC1_SIGNALING | MSigDb.Hallmark |
| E2F_TARGETS | MSigDb.Hallmark |
| MYC_TARGETS_V1 | MSigDb.Hallmark |
| MYC_TARGETS_V2 | MSigDb.Hallmark |
| EMT | MSigDb.Hallmark |
| INFLAMMATORY_RESPONSE | MSigDb.Hallmark |
| XENOBIOTIC_METABOLISM | MSigDb.Hallmark |
| FATTY_ACID_METABOLISM | MSigDb.Hallmark |
| OxPhos | MSigDb.Hallmark |
| GLYCOLYSIS | MSigDb.Hallmark |
| ROS | MSigDb.Hallmark |
| P53_PATHWAY | MSigDb.Hallmark |

| | |
|---|---|
| UV_RESPONSE_UP | MSigDb.Hallmark |
| UV_RESPONSE_DN | MSigDb.Hallmark |
| ANGIOGENESIS | MSigDb.Hallmark |
| HEME_METABOLISM | MSigDb.Hallmark |
| COAGULATION | MSigDb.Hallmark |
| IL2_STAT5_SIGNALING | MSigDb.Hallmark |
| BILE_ACID_METABOLISM | MSigDb.Hallmark |
| PEROXISOME | MSigDb.Hallmark |
| ALLOGRAFT_REJECTION | MSigDb.Hallmark |
| KRAS_SIGNALING_UP | MSigDb.Hallmark |
| KRAS_SIGNALING_DN | MSigDb.Hallmark |
| WNT_SIGNALLING | MSigDb.alternative.WNT.pathway// https://www.gsea-msigdb.org/gsea/msigdb/geneset_page.jsp?geneSetName=WNT_SIGNALING |
| FOETAL_INTESTINAL_UP | KimJensen.old.microarray// https://www.ebi.ac.uk/biostudies/arrayexpress/studies/E-MTAB-5246 |
| INTESTINAL_STEM_CELL | Merlos-Suárez, Anna, et al. Cell stem cell 8.5 (2011) |
| FA_vs_AF_UP | Chan et al. 2023 (this manuscript) |
| FOETAL_INTESTINAL_DN | KimJensen.old.microarray// https://www.ebi.ac.uk/biostudies/arrayexpress/studies/E-MTAB-5246 |
| FA_vs_AF_DN | Chan et al. 2023 (this manuscript) |
| YAP_COLON | Gregorieff, Alex, et al. Nature 526.7575 (2015) |
| Mustata_FOETAL_INTESTINAL | Mustata, Roxana C., et al. Cell reports 5.2 (2013) |
| Vallone_FOETAL_INTESTINAL | Fernandez Vallone, Valeria, et al. Development 143.9 (2016) |
| CBC_COLON | Vasquez, Ester Gil, et al. Cell stem cell 29.8 (2022) |
| RSC_COLON | Vasquez, Ester Gil, et al. Cell stem cell 29.8 (2022) |
| AbSC_COLON | Bala, Pratyusha, et al. Science Advances 9.13 (2023) |
| HIPPO_Reactome | https://www.gsea-msigdb.org/gsea/msigdb/human/geneset/REACTOME_SIGNALING_BY_HIPPO.html |
| HIPPO_REG_WikiP | https://www.gsea-msigdb.org/gsea/msigdb/human/geneset/WP_HIPPO_SIGNALING_REGULATION_PATHWAYS.html |
| HIPPO_WikiP | https://www.gsea-msigdb.org/gsea/msigdb/human/geneset/WP_HIPPOYAP_SIGNALING.html |
| YAP_MECHANOREG_WikiP | https://www.gsea-msigdb.org/gsea/msigdb/human/geneset/WP_MECHANOREGULATION_AND_PATHOLOGY_OF_YAPT ....html |
| Jensen2023_FOETAL_INTESTINAL | Pikkupeura et al. Science Advances. Jul 14;9(28) (2023) |
| Tape2023_CSC | ChrisTape.older.ver. (Biorxiv Preprint) |
| Tape2023_proCSC | ChrisTape.older.ver. (Biorxiv Preprint) |
| Tape2023_revCSC | ChrisTape.older.ver. (Biorxiv Preprint) |
| Tape2023_proCSC.v2 | Qin et al. Cell. Dec 7;186(25) (2023) |
| Tape2023_revCSC.v2 | Qin et al. Cell. Dec 7;186(25) (2023) |

**Extended Data Table 4 Table of de novo transcription factor identification results in F, A and W organoids**

| db internal ID | pwm id | TF | similarity to de novo motif | de novo motif | Comparison |
|---|---|---|---|---|---|
| 561 | MA0471.2 | E2F6 | 0.5293748 | motif1.FvsWT | FBXW7 vs WT |
| 318 | MA1122.1 | TFDP1 | 0.5138375 | motif1.FvsWT | FBXW7 vs WT |
| 632 | MA0528.2 | ZNF263 | 0.4843141 | motif1.FvsWT | FBXW7 vs WT |
| 627 | MA0814.2 | TFAP2C(var.2) | 0.4275616 | motif1.FvsWT | FBXW7 vs WT |
| 560 | MA1102.2 | CTCFL | 0.4034122 | motif1.FvsWT | FBXW7 vs WT |
| 20 | MA0130.1 | ZNF354C | 0.5568501 | motif4.FvsWT | FBXW7 vs WT |
| 402 | MA1514.1 | KLF17 | 0.4740363 | motif4.FvsWT | FBXW7 vs WT |
| 570 | MA0765.2 | ETV5 | 0.5906428 | motif2.FvsWT | FBXW7 vs WT |
| 566 | MA0640.2 | ELF3 | 0.5778408 | motif2.FvsWT | FBXW7 vs WT |
| 576 | MA0062.3 | GABPA | 0.5654622 | motif2.FvsWT | FBXW7 vs WT |
| 564 | MA0598.3 | EHF | 0.5322319 | motif2.FvsWT | FBXW7 vs WT |
| 46 | MA0076.2 | ELK4 | 0.5062427 | motif2.FvsWT | FBXW7 vs WT |
| 619 | MA0081.2 | SPIB | 0.4987504 | motif2.FvsWT | FBXW7 vs WT |
| 317 | MA1121.1 | TEAD2 | 0.4689957 | motif2.FvsWT | FBXW7 vs WT |
| 619 | MA0081.2 | SPIB | 0.5195268 | motif3.FvsWT | FBXW7 vs WT |
| 548 | MA1653.1 | ZNF148 | 0.5009792 | motif3.FvsWT | FBXW7 vs WT |
| 610 | MA0508.3 | PRDM1 | 0.4814322 | motif3.FvsWT | FBXW7 vs WT |
| 566 | MA0640.2 | ELF3 | 0.4405455 | motif3.FvsWT | FBXW7 vs WT |
| 24 | MA0152.1 | NFATC2 | 0.4330529 | motif3.FvsWT | FBXW7 vs WT |
| 564 | MA0598.3 | EHF | 0.4244305 | motif3.FvsWT | FBXW7 vs WT |
| 57 | MA0599.1 | KLF5 | 0.4242461 | motif3.FvsWT | FBXW7 vs WT |
| 449 | MA1564.1 | SP9 | 0.4220211 | motif3.FvsWT | FBXW7 vs WT |
| 576 | MA0062.3 | GABPA | 0.4157318 | motif3.FvsWT | FBXW7 vs WT |
| 82 | MA0648.1 | GSC | 0.4097036 | motif3.FvsWT | FBXW7 vs WT |
| 317 | MA1121.1 | TEAD2 | 0.7045162 | motif5.AvsFoe | APC vs Foetal |
| 625 | MA0809.2 | TEAD4 | 0.6987761 | motif5.AvsFoe | APC vs Foetal |
| 624 | MA0090.3 | TEAD1 | 0.6810789 | motif5.AvsFoe | APC vs Foetal |
| 47 | MA0258.2 | ESR2 | 0.4591278 | motif7.AvsFoe | APC vs Foetal |
| 538 | MA1643.1 | NFIB | 0.4577682 | motif7.AvsFoe | APC vs Foetal |
| 309 | MA1114.1 | PBX3 | 0.4264716 | motif7.AvsFoe | APC vs Foetal |
| 19 | MA0119.1 | NFIC::TLX1 | 0.4224025 | motif7.AvsFoe | APC vs Foetal |
| 578 | MA0482.2 | GATA4 | 0.5045814 | motif1.AvsFoe | APC vs Foetal |
| 352 | MA1152.1 | SOX15 | 0.4678806 | motif1.AvsFoe | APC vs Foetal |
| 619 | MA0081.2 | SPIB | 0.4659275 | motif1.AvsFoe | APC vs Foetal |
| 610 | MA0508.3 | PRDM1 | 0.465365 | motif1.AvsFoe | APC vs Foetal |
| 298 | MA0036.3 | GATA2 | 0.4648409 | motif1.AvsFoe | APC vs Foetal |
| 24 | MA0152.1 | NFATC2 | 0.4598157 | motif1.AvsFoe | APC vs Foetal |
| 579 | MA1104.2 | GATA6 | 0.4436686 | motif1.AvsFoe | APC vs Foetal |

| | | | | | |
|---|---|---|---|---|---|
| 566 | MA0640.2 | ELF3 | 0.4223688 | motif1.AvsFoe | APC vs Foetal |
| 61 | MA0625.1 | NFATC3 | 0.4170524 | motif1.AvsFoe | APC vs Foetal |
| 577 | MA0035.4 | GATA1 | 0.4144784 | motif1.AvsFoe | APC vs Foetal |
| 576 | MA0062.3 | GABPA | 0.6569089 | motif3.AvsFoe | APC vs Foetal |
| 564 | MA0598.3 | EHF | 0.6561789 | motif3.AvsFoe | APC vs Foetal |
| 619 | MA0081.2 | SPIB | 0.6239054 | motif3.AvsFoe | APC vs Foetal |
| 48 | MA0050.2 | IRF1 | 0.5130528 | motif3.AvsFoe | APC vs Foetal |
| 551 | MA1656.1 | ZNF449 | 0.5547182 | motif4.AvsFoe | APC vs Foetal |
| 478 | MA1599.1 | ZNF682 | 0.511596 | motif4.AvsFoe | APC vs Foetal |
| 587 | MA0039.4 | KLF4 | 0.4879408 | motif4.AvsFoe | APC vs Foetal |
| 521 | MA0516.2 | SP2 | 0.485105 | motif4.AvsFoe | APC vs Foetal |
| 169 | MA0741.1 | KLF16 | 0.4685539 | motif4.AvsFoe | APC vs Foetal |
| 563 | MA0162.4 | EGR1 | 0.4670903 | motif4.AvsFoe | APC vs Foetal |
| 47 | MA0258.2 | ESR2 | 0.4613923 | motif4.AvsFoe | APC vs Foetal |
| 163 | MA0733.1 | EGR4 | 0.4586193 | motif4.AvsFoe | APC vs Foetal |
| 522 | MA0746.2 | SP3 | 0.4536906 | motif4.AvsFoe | APC vs Foetal |
| 401 | MA1513.1 | KLF15 | 0.4477592 | motif4.AvsFoe | APC vs Foetal |
| 403 | MA1515.1 | KLF2 | 0.4439192 | motif4.AvsFoe | APC vs Foetal |
| 400 | MA1512.1 | KLF11 | 0.4409988 | motif4.AvsFoe | APC vs Foetal |
| 548 | MA1653.1 | ZNF148 | 0.4393953 | motif4.AvsFoe | APC vs Foetal |
| 479 | MA1600.1 | ZNF684 | 0.4359734 | motif4.AvsFoe | APC vs Foetal |
| 566 | MA0640.2 | ELF3 | 0.4595839 | motif2.AvsFoe | APC vs Foetal |
| 57 | MA0599.1 | KLF5 | 0.4510012 | motif2.AvsFoe | APC vs Foetal |
| 193 | MA0498.2 | MEIS1 | 0.4181185 | motif2.AvsFoe | APC vs Foetal |
| 564 | MA0598.3 | EHF | 0.4172832 | motif2.AvsFoe | APC vs Foetal |
| 476 | MA1596.1 | ZNF460 | 0.4156459 | motif2.AvsFoe | APC vs Foetal |
| 20 | MA0130.1 | ZNF354C | 0.4110677 | motif2.AvsFoe | APC vs Foetal |
| 576 | MA0062.3 | GABPA | 0.4031677 | motif2.AvsFoe | APC vs Foetal |
| 565 | MA0473.3 | ELF1 | 0.6256227 | motif2.AvsWT | APC vs WT |
| 319 | MA0750.2 | ZBTB7A | 0.5932214 | motif2.AvsWT | APC vs WT |
| 397 | MA1508.1 | IKZF1 | 0.5757949 | motif2.AvsWT | APC vs WT |
| 477 | MA1597.1 | ZNF528 | 0.5586696 | motif2.AvsWT | APC vs WT |
| 568 | MA0761.2 | ETV1 | 0.5584296 | motif2.AvsWT | APC vs WT |
| 71 | MA0136.2 | ELF5 | 0.5541939 | motif2.AvsWT | APC vs WT |
| 70 | MA0641.1 | ELF4 | 0.5314541 | motif2.AvsWT | APC vs WT |
| 49 | MA0137.3 | STAT1 | 0.5251467 | motif2.AvsWT | APC vs WT |
| 22 | MA0149.1 | EWSR1-FLI1 | 0.5114498 | motif2.AvsWT | APC vs WT |
| 569 | MA0764.2 | ETV4 | 0.5064976 | motif2.AvsWT | APC vs WT |
| 48 | MA0050.2 | IRF1 | 0.5009438 | motif3.AvsWT | APC vs WT |
| 610 | MA0508.3 | PRDM1 | 0.438467 | motif3.AvsWT | APC vs WT |
| 352 | MA1152.1 | SOX15 | 0.426994 | motif3.AvsWT | APC vs WT |
| 61 | MA0625.1 | NFATC3 | 0.4178249 | motif3.AvsWT | APC vs WT |
| 619 | MA0081.2 | SPIB | 0.4073858 | motif3.AvsWT | APC vs WT |
| 20 | MA0130.1 | ZNF354C | 0.5726547 | motif5.AvsWT | APC vs WT |

| | | | | | |
|---|---|---|---|---|---|
| 402 | MA1514.1 | KLF17 | 0.4764035 | motif5.AvsWT | APC vs WT |
| 618 | MA0080.5 | SPI1 | 0.5022072 | motif4.AvsWT | APC vs WT |
| 450 | MA1565.1 | TBX18 | 0.4851093 | motif4.AvsWT | APC vs WT |
| 547 | MA1652.1 | ZKSCAN5 | 0.4812596 | motif4.AvsWT | APC vs WT |
| 632 | MA0528.2 | ZNF263 | 0.4601932 | motif4.AvsWT | APC vs WT |
| 35 | MA0489.1 | JUN(var.2) | 0.4331843 | motif4.AvsWT | APC vs WT |
| 22 | MA0149.1 | EWSR1-FLI1 | 0.4252739 | motif4.AvsWT | APC vs WT |
| 4 | MA0057.1 | MZF1(var.2) | 0.4171988 | motif4.AvsWT | APC vs WT |
| 401 | MA1513.1 | KLF15 | 0.6561247 | motif1.AvsWT | APC vs WT |
| 587 | MA0039.4 | KLF4 | 0.6238827 | motif1.AvsWT | APC vs WT |
| 548 | MA1653.1 | ZNF148 | 0.6154464 | motif1.AvsWT | APC vs WT |
| 169 | MA0741.1 | KLF16 | 0.6000897 | motif1.AvsWT | APC vs WT |
| 410 | MA1522.1 | MAZ | 0.589003 | motif1.AvsWT | APC vs WT |
| 449 | MA1564.1 | SP9 | 0.581299 | motif1.AvsWT | APC vs WT |
| 400 | MA1512.1 | KLF11 | 0.5766117 | motif1.AvsWT | APC vs WT |
| 545 | MA1650.1 | ZBTB14 | 0.5474092 | motif1.AvsWT | APC vs WT |
| 57 | MA0599.1 | KLF5 | 0.5389704 | motif1.AvsWT | APC vs WT |
| 399 | MA1511.1 | KLF10 | 0.5330812 | motif1.AvsWT | APC vs WT |
| 522 | MA0746.2 | SP3 | 0.5310723 | motif1.AvsWT | APC vs WT |
| 170 | MA0747.1 | SP8 | 0.5290076 | motif1.AvsWT | APC vs WT |
| 403 | MA1515.1 | KLF2 | 0.5210434 | motif1.AvsWT | APC vs WT |
| 588 | MA1107.2 | KLF9 | 0.5154122 | motif1.AvsWT | APC vs WT |
| 61 | MA0625.1 | NFATC3 | 0.4768709 | motif3.FoevsWT | Foetal vs WT |
| 56 | MA0597.1 | THAP1 | 0.452531 | motif3.FoevsWT | Foetal vs WT |
| 528 | MA1633.1 | BACH1 | 0.6959529 | motif4.FoevsWT | Foetal vs WT |
| 592 | MA0496.3 | MAFK | 0.680554 | motif4.FoevsWT | Foetal vs WT |
| 338 | MA1137.1 | FOSL1::JUNB | 0.6701658 | motif4.FoevsWT | Foetal vs WT |
| 599 | MA0089.2 | NFE2L1 | 0.6279242 | motif4.FoevsWT | Foetal vs WT |
| 571 | MA0477.2 | FOSL1 | 0.6260315 | motif4.FoevsWT | Foetal vs WT |
| 35 | MA0489.1 | JUN(var.2) | 0.6234823 | motif4.FoevsWT | Foetal vs WT |
| 586 | MA0491.2 | JUND | 0.622893 | motif4.FoevsWT | Foetal vs WT |
| 38 | MA0501.1 | MAF::NFE2 | 0.6162797 | motif4.FoevsWT | Foetal vs WT |
| 409 | MA1521.1 | MAFA | 0.6070557 | motif4.FoevsWT | Foetal vs WT |
| 342 | MA1142.1 | FOSL1::JUND | 0.606453 | motif4.FoevsWT | Foetal vs WT |
| 585 | MA0490.2 | JUNB | 0.6052837 | motif4.FoevsWT | Foetal vs WT |
| 408 | MA1520.1 | MAF | 0.5974487 | motif4.FoevsWT | Foetal vs WT |
| 401 | MA1513.1 | KLF15 | 0.6536718 | motif1.FoevsWT | Foetal vs WT |
| 587 | MA0039.4 | KLF4 | 0.6274664 | motif1.FoevsWT | Foetal vs WT |
| 548 | MA1653.1 | ZNF148 | 0.6108673 | motif1.FoevsWT | Foetal vs WT |
| 169 | MA0741.1 | KLF16 | 0.601685 | motif1.FoevsWT | Foetal vs WT |
| 410 | MA1522.1 | MAZ | 0.5798422 | motif1.FoevsWT | Foetal vs WT |
| 400 | MA1512.1 | KLF11 | 0.5785008 | motif1.FoevsWT | Foetal vs WT |
| 545 | MA1650.1 | ZBTB14 | 0.5526084 | motif1.FoevsWT | Foetal vs WT |
| 563 | MA0162.4 | EGR1 | 0.5515962 | motif1.FoevsWT | Foetal vs WT |

| | | | | | |
|---|---|---|---|---|---|
| 403 | MA1515.1 | KLF2 | 0.533581 | motif1.FoevsWT | Foetal vs WT |
| 57 | MA0599.1 | KLF5 | 0.5333037 | motif1.FoevsWT | Foetal vs WT |
| 399 | MA1511.1 | KLF10 | 0.533099 | motif1.FoevsWT | Foetal vs WT |
| 405 | MA1517.1 | KLF6 | 0.5099714 | motif1.FoevsWT | Foetal vs WT |
| 561 | MA0471.2 | E2F6 | 0.5282559 | motif2.FoevsWT | Foetal vs WT |
| 632 | MA0528.2 | ZNF263 | 0.5140558 | motif2.FoevsWT | Foetal vs WT |
| 547 | MA1652.1 | ZKSCAN5 | 0.5132787 | motif2.FoevsWT | Foetal vs WT |
| 397 | MA1508.1 | IKZF1 | 0.4475741 | motif2.FoevsWT | Foetal vs WT |
| 4 | MA0057.1 | MZF1(var.2) | 0.4361988 | motif2.FoevsWT | Foetal vs WT |
| 565 | MA0473.3 | ELF1 | 0.434381 | motif2.FoevsWT | Foetal vs WT |
| 518 | MA0867.2 | SOX4 | 0.4236869 | motif2.FoevsWT | Foetal vs WT |
| 546 | MA1651.1 | ZFP42 | 0.4725357 | motif7.FvsFoe | FBXW7 vs Foetal |
| 42 | MA0513.1 | SMAD2::SMAD3::SMAD4 | 0.4417832 | motif7.FvsFoe | FBXW7 vs Foetal |
| 402 | MA1514.1 | KLF17 | 0.4138352 | motif7.FvsFoe | FBXW7 vs Foetal |
| 476 | MA1596.1 | ZNF460 | 0.4091909 | motif7.FvsFoe | FBXW7 vs Foetal |
| 548 | MA1653.1 | ZNF148 | 0.4072226 | motif7.FvsFoe | FBXW7 vs Foetal |
| 416 | MA1529.1 | NHLH2 | 0.4043787 | motif7.FvsFoe | FBXW7 vs Foetal |
| 338 | MA1137.1 | FOSL1::JUNB | 0.6290006 | motif10.FvsFoe | FBXW7 vs Foetal |
| 586 | MA0491.2 | JUND | 0.6081693 | motif10.FvsFoe | FBXW7 vs Foetal |
| 571 | MA0477.2 | FOSL1 | 0.6012091 | motif10.FvsFoe | FBXW7 vs Foetal |
| 528 | MA1633.1 | BACH1 | 0.5995408 | motif10.FvsFoe | FBXW7 vs Foetal |
| 592 | MA0496.3 | MAFK | 0.5976529 | motif10.FvsFoe | FBXW7 vs Foetal |
| 529 | MA1634.1 | BATF | 0.5964315 | motif10.FvsFoe | FBXW7 vs Foetal |
| 485 | MA0462.2 | BATF::JUN | 0.5925071 | motif10.FvsFoe | FBXW7 vs Foetal |
| 585 | MA0490.2 | JUNB | 0.591946 | motif10.FvsFoe | FBXW7 vs Foetal |
| 35 | MA0489.1 | JUN(var.2) | 0.5809788 | motif10.FvsFoe | FBXW7 vs Foetal |
| 599 | MA0089.2 | NFE2L1 | 0.5725839 | motif10.FvsFoe | FBXW7 vs Foetal |
| 409 | MA1521.1 | MAFA | 0.5585196 | motif10.FvsFoe | FBXW7 vs Foetal |
| 555 | MA0835.2 | BATF3 | 0.5571789 | motif10.FvsFoe | FBXW7 vs Foetal |
| 408 | MA1520.1 | MAF | 0.5562729 | motif10.FvsFoe | FBXW7 vs Foetal |
| 32 | MA0478.1 | FOSL2 | 0.546946 | motif10.FvsFoe | FBXW7 vs Foetal |
| 335 | MA1134.1 | FOS::JUNB | 0.5450862 | motif10.FvsFoe | FBXW7 vs Foetal |
| 342 | MA1142.1 | FOSL1::JUND | 0.5391123 | motif10.FvsFoe | FBXW7 vs Foetal |
| 331 | MA1130.1 | FOSL2::JUN | 0.5301662 | motif10.FvsFoe | FBXW7 vs Foetal |
| 333 | MA1132.1 | JUN::JUNB | 0.52869 | motif10.FvsFoe | FBXW7 vs Foetal |
| 344 | MA1144.1 | FOSL2::JUND | 0.5215272 | motif10.FvsFoe | FBXW7 vs Foetal |
| 329 | MA1128.1 | FOSL1::JUN | 0.5178963 | motif10.FvsFoe | FBXW7 vs Foetal |
| 339 | MA1138.1 | FOSL2::JUNB | 0.5148668 | motif10.FvsFoe | FBXW7 vs Foetal |
| 92 | MA0655.1 | JDP2 | 0.511349 | motif10.FvsFoe | FBXW7 vs Foetal |
| 31 | MA0476.1 | FOS | 0.5112956 | motif10.FvsFoe | FBXW7 vs Foetal |
| 592 | MA0496.3 | MAFK | 0.462806 | motif5.FvsFoe | FBXW7 vs Foetal |
| 309 | MA1114.1 | PBX3 | 0.4175988 | motif5.FvsFoe | FBXW7 vs Foetal |
| 565 | MA0473.3 | ELF1 | 0.5874015 | motif3.FvsFoe | FBXW7 vs Foetal |
| 70 | MA0641.1 | ELF4 | 0.5600106 | motif3.FvsFoe | FBXW7 vs Foetal |

| | | | | | |
|---|---|---|---|---|---|
| 71 | MA0136.2 | ELF5 | 0.5554768 | motif3.FvsFoe | FBXW7 vs Foetal |
| 397 | MA1508.1 | IKZF1 | 0.5478055 | motif3.FvsFoe | FBXW7 vs Foetal |
| 376 | MA1483.1 | ELF2 | 0.5242166 | motif3.FvsFoe | FBXW7 vs Foetal |
| 477 | MA1597.1 | ZNF528 | 0.5016066 | motif3.FvsFoe | FBXW7 vs Foetal |
| 118 | MA0686.1 | SPDEF | 0.4891945 | motif3.FvsFoe | FBXW7 vs Foetal |
| 568 | MA0761.2 | ETV1 | 0.4864729 | motif3.FvsFoe | FBXW7 vs Foetal |
| 464 | MA1579.1 | ZBTB26 | 0.4826854 | motif3.FvsFoe | FBXW7 vs Foetal |
| 49 | MA0137.3 | STAT1 | 0.4799056 | motif3.FvsFoe | FBXW7 vs Foetal |
| 319 | MA0750.2 | ZBTB7A | 0.4658223 | motif3.FvsFoe | FBXW7 vs Foetal |
| 632 | MA0528.2 | ZNF263 | 0.6086341 | motif1.FvsFoe | FBXW7 vs Foetal |
| 618 | MA0080.5 | SPI1 | 0.5579887 | motif1.FvsFoe | FBXW7 vs Foetal |
| 561 | MA0471.2 | E2F6 | 0.5316274 | motif1.FvsFoe | FBXW7 vs Foetal |
| 397 | MA1508.1 | IKZF1 | 0.5167911 | motif1.FvsFoe | FBXW7 vs Foetal |
| 565 | MA0473.3 | ELF1 | 0.5011948 | motif1.FvsFoe | FBXW7 vs Foetal |
| 22 | MA0149.1 | EWSR1-FLI1 | 0.4958431 | motif1.FvsFoe | FBXW7 vs Foetal |
| 568 | MA0761.2 | ETV1 | 0.4870791 | motif1.FvsFoe | FBXW7 vs Foetal |
| 587 | MA0039.4 | KLF4 | 0.7034798 | motif9.FvsFoe | FBXW7 vs Foetal |
| 57 | MA0599.1 | KLF5 | 0.6214743 | motif9.FvsFoe | FBXW7 vs Foetal |
| 588 | MA1107.2 | KLF9 | 0.6192501 | motif9.FvsFoe | FBXW7 vs Foetal |
| 400 | MA1512.1 | KLF11 | 0.6018859 | motif9.FvsFoe | FBXW7 vs Foetal |
| 522 | MA0746.2 | SP3 | 0.593456 | motif9.FvsFoe | FBXW7 vs Foetal |
| 405 | MA1517.1 | KLF6 | 0.5832665 | motif9.FvsFoe | FBXW7 vs Foetal |
| 399 | MA1511.1 | KLF10 | 0.5712182 | motif9.FvsFoe | FBXW7 vs Foetal |
| 618 | MA0080.5 | SPI1 | 0.5541379 | motif6.FvsFoe | FBXW7 vs Foetal |
| 315 | MA0442.2 | SOX10 | 0.5068245 | motif6.FvsFoe | FBXW7 vs Foetal |
| 44 | MA0523.1 | TCF7L2 | 0.4923421 | motif6.FvsFoe | FBXW7 vs Foetal |
| 322 | MA1125.1 | ZNF384 | 0.4839554 | motif6.FvsFoe | FBXW7 vs Foetal |
| 617 | MA0143.4 | SOX2 | 0.4779196 | motif6.FvsFoe | FBXW7 vs Foetal |
| 53 | MA0593.1 | FOXP2 | 0.4634188 | motif6.FvsFoe | FBXW7 vs Foetal |
| 556 | MA0465.2 | CDX2 | 0.4526045 | motif6.FvsFoe | FBXW7 vs Foetal |
| 604 | MA0679.2 | ONECUT1 | 0.4520453 | motif6.FvsFoe | FBXW7 vs Foetal |
| 518 | MA0867.2 | SOX4 | 0.4411986 | motif6.FvsFoe | FBXW7 vs Foetal |
| 178 | MA0757.1 | ONECUT3 | 0.4169856 | motif6.FvsFoe | FBXW7 vs Foetal |
| 11 | MA0073.1 | RREB1 | 0.5109962 | motif4.FvsFoe | FBXW7 vs Foetal |
| 221 | MA0511.2 | RUNX2 | 0.4987379 | motif4.FvsFoe | FBXW7 vs Foetal |
| 588 | MA1107.2 | KLF9 | 0.4650438 | motif4.FvsFoe | FBXW7 vs Foetal |
| 613 | MA0684.2 | RUNX3 | 0.4464971 | motif4.FvsFoe | FBXW7 vs Foetal |
| 129 | MA0695.1 | ZBTB7C | 0.4377965 | motif4.FvsFoe | FBXW7 vs Foetal |
| 20 | MA0130.1 | ZNF354C | 0.4289784 | motif4.FvsFoe | FBXW7 vs Foetal |
| 25 | MA0155.1 | INSM1 | 0.527211 | motif2.FvsFoe | FBXW7 vs Foetal |
| 627 | MA0814.2 | TFAP2C(var.2) | 0.4904893 | motif2.FvsFoe | FBXW7 vs Foetal |
| 267 | MA0106.3 | TP53 | 0.4440641 | motif2.FvsFoe | FBXW7 vs Foetal |
| 28 | MA0163.1 | PLAG1 | 0.4415809 | motif2.FvsFoe | FBXW7 vs Foetal |
| 268 | MA0861.1 | TP73 | 0.4384959 | motif2.FvsFoe | FBXW7 vs Foetal |

| | | | | | |
|---|---|---|---|---|---|
| 626 | MA0003.4 | TFAP2A | 0.4383031 | motif2.FvsFoe | FBXW7 vs Foetal |
| 551 | MA1656.1 | ZNF449 | 0.4536013 | motif8.FvsFoe | FBXW7 vs Foetal |
| 397 | MA1508.1 | IKZF1 | 0.6194757 | motif8.FvsA | FBXW7 vs APC |
| 569 | MA0764.2 | ETV4 | 0.590983 | motif8.FvsA | FBXW7 vs APC |
| 568 | MA0761.2 | ETV1 | 0.585426 | motif8.FvsA | FBXW7 vs APC |
| 565 | MA0473.3 | ELF1 | 0.582969 | motif8.FvsA | FBXW7 vs APC |
| 319 | MA0750.2 | ZBTB7A | 0.5719968 | motif8.FvsA | FBXW7 vs APC |
| 22 | MA0149.1 | EWSR1-FLI1 | 0.554565 | motif8.FvsA | FBXW7 vs APC |
| 377 | MA1484.1 | ETS2 | 0.5395569 | motif8.FvsA | FBXW7 vs APC |
| 70 | MA0641.1 | ELF4 | 0.5370904 | motif8.FvsA | FBXW7 vs APC |
| 618 | MA0080.5 | SPI1 | 0.5196112 | motif8.FvsA | FBXW7 vs APC |
| 547 | MA1652.1 | ZKSCAN5 | 0.5131197 | motif8.FvsA | FBXW7 vs APC |
| 182 | MA0474.2 | ERG | 0.5072231 | motif8.FvsA | FBXW7 vs APC |
| 184 | MA0762.1 | ETV2 | 0.5044169 | motif8.FvsA | FBXW7 vs APC |
| 71 | MA0136.2 | ELF5 | 0.5033707 | motif8.FvsA | FBXW7 vs APC |
| 183 | MA0098.3 | ETS1 | 0.5027285 | motif8.FvsA | FBXW7 vs APC |
| 186 | MA0156.2 | FEV | 0.4959729 | motif8.FvsA | FBXW7 vs APC |
| 76 | MA0475.2 | FLI1 | 0.4950672 | motif8.FvsA | FBXW7 vs APC |
| 181 | MA0760.1 | ERF | 0.4927738 | motif8.FvsA | FBXW7 vs APC |
| 476 | MA1596.1 | ZNF460 | 0.536358 | motif5.FvsA | FBXW7 vs APC |
| 546 | MA1651.1 | ZFP42 | 0.4559162 | motif5.FvsA | FBXW7 vs APC |
| 320 | MA0103.3 | ZEB1 | 0.4312871 | motif5.FvsA | FBXW7 vs APC |
| 548 | MA1653.1 | ZNF148 | 0.4261082 | motif5.FvsA | FBXW7 vs APC |
| 221 | MA0511.2 | RUNX2 | 0.5494413 | motif4.FvsA | FBXW7 vs APC |
| 613 | MA0684.2 | RUNX3 | 0.4857266 | motif4.FvsA | FBXW7 vs APC |
| 20 | MA0130.1 | ZNF354C | 0.4401673 | motif4.FvsA | FBXW7 vs APC |
| 592 | MA0496.3 | MAFK | 0.617434 | motif7.FvsA | FBXW7 vs APC |
| 409 | MA1521.1 | MAFA | 0.5904365 | motif7.FvsA | FBXW7 vs APC |
| 408 | MA1520.1 | MAF | 0.5757799 | motif7.FvsA | FBXW7 vs APC |
| 338 | MA1137.1 | FOSL1::JUNB | 0.5730276 | motif7.FvsA | FBXW7 vs APC |
| 336 | MA1135.1 | FOSB::JUNB | 0.5642777 | motif7.FvsA | FBXW7 vs APC |
| 335 | MA1134.1 | FOS::JUNB | 0.5631371 | motif7.FvsA | FBXW7 vs APC |
| 333 | MA1132.1 | JUN::JUNB | 0.555136 | motif7.FvsA | FBXW7 vs APC |
| 528 | MA1633.1 | BACH1 | 0.5521391 | motif7.FvsA | FBXW7 vs APC |
| 344 | MA1144.1 | FOSL2::JUND | 0.551166 | motif7.FvsA | FBXW7 vs APC |
| 339 | MA1138.1 | FOSL2::JUNB | 0.5490041 | motif7.FvsA | FBXW7 vs APC |
| 342 | MA1142.1 | FOSL1::JUND | 0.5461288 | motif7.FvsA | FBXW7 vs APC |
| 92 | MA0655.1 | JDP2 | 0.5342542 | motif7.FvsA | FBXW7 vs APC |
| 329 | MA1128.1 | FOSL1::JUN | 0.5224258 | motif7.FvsA | FBXW7 vs APC |
| 586 | MA0491.2 | JUND | 0.521567 | motif7.FvsA | FBXW7 vs APC |
| 571 | MA0477.2 | FOSL1 | 0.5213872 | motif7.FvsA | FBXW7 vs APC |
| 331 | MA1130.1 | FOSL2::JUN | 0.5205519 | motif7.FvsA | FBXW7 vs APC |
| 31 | MA0476.1 | FOS | 0.5190281 | motif7.FvsA | FBXW7 vs APC |
| 341 | MA1141.1 | FOS::JUND | 0.5041699 | motif7.FvsA | FBXW7 vs APC |

| | | | | | |
|---|---|---|---|---|---|
| 35 | MA0489.1 | JUN(var.2) | 0.503129 | motif7.FvsA | FBXW7 vs APC |
| 195 | MA0775.1 | MEIS3 | 0.4408985 | motif6.FvsA | FBXW7 vs APC |
| 561 | MA0471.2 | E2F6 | 0.4234415 | motif6.FvsA | FBXW7 vs APC |
| 202 | MA0781.1 | PAX9 | 0.4181158 | motif6.FvsA | FBXW7 vs APC |
| 477 | MA1597.1 | ZNF528 | 0.4100592 | motif6.FvsA | FBXW7 vs APC |
| 39 | MA0504.1 | NR2C2 | 0.4058812 | motif6.FvsA | FBXW7 vs APC |
| 200 | MA0779.1 | PAX1 | 0.4035235 | motif6.FvsA | FBXW7 vs APC |
| 561 | MA0471.2 | E2F6 | 0.4884101 | motif1.FvsA | FBXW7 vs APC |
| 632 | MA0528.2 | ZNF263 | 0.6197193 | motif2.FvsA | FBXW7 vs APC |
| 561 | MA0471.2 | E2F6 | 0.5440533 | motif2.FvsA | FBXW7 vs APC |
| 560 | MA1102.2 | CTCFL | 0.503502 | motif3.FvsA | FBXW7 vs APC |
| 454 | MA1569.1 | TFAP2E | 0.4823678 | motif3.FvsA | FBXW7 vs APC |
| 626 | MA0003.4 | TFAP2A | 0.4707305 | motif3.FvsA | FBXW7 vs APC |
| 627 | MA0814.2 | TFAP2C(var.2) | 0.4670565 | motif3.FvsA | FBXW7 vs APC |
| 561 | MA0471.2 | E2F6 | 0.4591359 | motif3.FvsA | FBXW7 vs APC |
| 302 | MA0104.4 | MYCN | 0.4509939 | motif3.FvsA | FBXW7 vs APC |
| 233 | MA0812.1 | TFAP2B(var.2) | 0.4426652 | motif3.FvsA | FBXW7 vs APC |
| 22 | MA0149.1 | EWSR1-FLI1 | 0.4292005 | motif3.FvsA | FBXW7 vs APC |
| 319 | MA0750.2 | ZBTB7A | 0.411418 | motif3.FvsA | FBXW7 vs APC |
| 21 | MA0139.1 | CTCF | 0.4032005 | motif3.FvsA | FBXW7 vs APC |
| 19 | MA0119.1 | NFIC::TLX1 | 0.4021992 | motif3.FvsA | FBXW7 vs APC |
| 303 | MA1109.1 | NEUROD1 | 0.4020819 | motif3.FvsA | FBXW7 vs APC |

Annotation of the de novo motifs is as per JASPAR 2020 release (R package)

**Extended Data Table 5 GSEA results for Hallmark and stem cell signatures on pre-operative biopsies from the FOxTROT study**

| pathway | pval | padj | log2err | ES | NES | size |
|---|---|---|---|---|---|---|
| FAvsAF.UP | 0.0001247 | 0.002276 | 0.5188481 | 0.3563436 | 1.6772558 | 195 |
| AFvsWT.DN | 0.000295 | 0.0043063 | 0.4984931 | 0.3271019 | 1.5848787 | 256 |
| ANGIOGENESIS | 0.0233539 | 0.1398085 | 0.3524879 | 0.4311433 | 1.5138647 | 35 |
| APICAL_SURFACE | 0.0361476 | 0.1398085 | 0.3217759 | 0.4117259 | 1.503241 | 40 |
| FAvsWT.UP | 0.0033316 | 0.028024 | 0.4317077 | 0.3259478 | 1.4823949 | 158 |
| PROTEIN_SECRETION | 0.031438 | 0.1398085 | 0.3217759 | 0.3322146 | 1.4223107 | 95 |
| Mustata_FOETAL_INTESTINAL | 0.0059998 | 0.0437985 | 0.4070179 | 0.2922654 | 1.4060529 | 239 |
| Jensen2023_FOETAL_INTESTINAL | 0.0370115 | 0.1398085 | 0.3217759 | 0.3023846 | 1.3655072 | 146 |
| FAvsWT.DN | 0.0360578 | 0.1398085 | 0.3217759 | 0.3029912 | 1.3586869 | 138 |
| Tape2023_proCSC.v2 | 0.0466761 | 0.162255 | 0.2489111 | 0.3157503 | 1.3577762 | 97 |
| EPITHELIAL_MESENCHYMAL_TRANSITION | 0.031113 | 0.1398085 | 0.3524879 | 0.2880912 | 1.3547987 | 192 |
| AvsWT.UP | 0.0277413 | 0.1398085 | 0.3524879 | 0.276496 | 1.3227687 | 220 |
| FvsWT.UP | 0.0383037 | 0.1398085 | 0.2712886 | 0.2897668 | 1.3166217 | 149 |
| E2F_TARGETS | 0.0349933 | 0.1398085 | 0.2820134 | 0.2786211 | 1.3077673 | 191 |
| INTESTINAL_STEM_CELL | 0.0755287 | 0.2297331 | 0.1999152 | 0.3263554 | 1.307085 | 65 |
| FvsWT.DN | 0.055788 | 0.1851147 | 0.2249661 | 0.2898481 | 1.304239 | 140 |
| Vallone_FOETAL_INTESTINAL | 0.0834532 | 0.2324841 | 0.1847065 | 0.2914605 | 1.2653242 | 108 |
| IL2_STAT5_SIGNALING | 0.0831099 | 0.2324841 | 0.1782199 | 0.264356 | 1.2350589 | 185 |
| AvsWT.DN | 0.0902873 | 0.2353918 | 0.1723243 | 0.2682899 | 1.2288677 | 161 |
| FATTY_ACID_METABOLISM | 0.0973937 | 0.2451634 | 0.1656567 | 0.2698092 | 1.2248552 | 153 |
| WNT_SIGNALLING | 0.12627 | 0.2793244 | 0.1482615 | 0.2940358 | 1.2218836 | 80 |
| TGF_BETA_SIGNALING | 0.1804281 | 0.3658682 | 0.1250334 | 0.3127649 | 1.2012814 | 52 |
| CBC_COLON | 0.1076923 | 0.2599779 | 0.1511488 | 0.2368958 | 1.1695527 | 318 |
| KRAS_SIGNALING_UP | 0.1879195 | 0.37076 | 0.1133129 | 0.2450923 | 1.1492738 | 189 |
| FOETAL_INTESTINAL_UP | 0.1625806 | 0.3390968 | 0.1204334 | 0.2333167 | 1.1393742 | 283 |
| YAP_MECHANOREG_WikiP | 0.259434 | 0.4557515 | 0.1028218 | 0.3103 | 1.1349225 | 41 |
| PEROXISOME | 0.2362869 | 0.4460136 | 0.1017139 | 0.2630528 | 1.1329107 | 99 |
| HIPPO_Reactome | 0.3065068 | 0.4894318 | 0.0978773 | 0.3691716 | 1.1222924 | 20 |
| HIPPO_REG_WikiP | 0.255102 | 0.4557515 | 0.0992333 | 0.2666788 | 1.1208441 | 87 |
| CHOLESTEROL_HOMEOSTASIS | 0.2664671 | 0.4557515 | 0.0982123 | 0.2771293 | 1.1164639 | 68 |
| HEDGEHOG_SIGNALING | 0.3170347 | 0.4894318 | 0.0911073 | 0.3086629 | 1.0929658 | 36 |
| COAGULATION | 0.3239832 | 0.4894318 | 0.0833634 | 0.2427588 | 1.0781768 | 123 |
| AFvsWT.UP | 0.3352273 | 0.4894318 | 0.0822055 | 0.2491241 | 1.0746079 | 101 |
| COMPLEMENT_INNATE_IMMUNE_SYSTEM | 0.3150134 | 0.4894318 | 0.0824344 | 0.2288814 | 1.0727907 | 188 |
| UV_RESPONSE_DN | 0.3323944 | 0.4894318 | 0.0822055 | 0.2357611 | 1.055091 | 135 |
| ADIPOGENESIS | 0.3624161 | 0.5087765 | 0.0751182 | 0.2243603 | 1.0520585 | 189 |
| GLYCOLYSIS | 0.3579088 | 0.5087765 | 0.0756946 | 0.2236658 | 1.0490874 | 190 |
| ESTROGEN_RESPONSE_EARLY | 0.4021448 | 0.5436402 | 0.0697793 | 0.2190827 | 1.0275906 | 190 |
| RSC_COLON | 0.4419226 | 0.5760776 | 0.0649408 | 0.215827 | 1.0149639 | 192 |

| | | | | | | |
|---|---|---|---|---|---|---|
| HYPOXIA | 0.5046729 | 0.6463355 | 0.0585938 | 0.2094731 | 0.9850837 | 192 |
| ANDROGEN_RESPONSE | 0.5190948 | 0.6533434 | 0.0599892 | 0.2270324 | 0.976275 | 97 |
| MYC_TARGETS_V1 | 0.5567423 | 0.6765916 | 0.0540088 | 0.204645 | 0.9623783 | 192 |
| AbSC_COLON | 0.6136681 | 0.7225447 | 0.0515309 | 0.2075366 | 0.9355151 | 145 |
| ESTROGEN_RESPONSE_LATE | 0.6340483 | 0.7232113 | 0.048215 | 0.1988476 | 0.9326794 | 190 |
| Tape2023_revCSC.v2 | 0.6479058 | 0.7265009 | 0.0461379 | 0.1950541 | 0.9303249 | 214 |
| G2M_CHECKPOINT | 0.6568365 | 0.7265009 | 0.0466015 | 0.1970612 | 0.9229863 | 187 |
| NOTCH_SIGNALING | 0.6256 | 0.7232113 | 0.057006 | 0.2621335 | 0.8968581 | 32 |
| APICAL_JUNCTION | 0.7127517 | 0.7540706 | 0.0429311 | 0.1912415 | 0.8947145 | 186 |
| YAP_COLON | 0.740988 | 0.7727446 | 0.0409056 | 0.1886829 | 0.8873141 | 192 |
| HIPPO_WikiP | 0.7043919 | 0.7540706 | 0.0540088 | 0.2627701 | 0.8152954 | 22 |
| KRAS_SIGNALING_DN | 0.9099591 | 0.9099591 | 0.0322878 | 0.1746991 | 0.8006373 | 163 |
| BILE_ACID_METABOLISM | 0.8987342 | 0.9099591 | 0.0343431 | 0.1798368 | 0.7745176 | 99 |
| REACTIVE_OXYGEN_SPECIES_PATHWAY | 0.8580442 | 0.8822144 | 0.0418239 | 0.2039663 | 0.7620324 | 44 |
| OXIDATIVE_PHOSPHORYLATION | 0.6914063 | 0.7533232 | 0.09855 | -0.180469 | -0.93555 | 183 |
| FOETAL_INTESTINAL_DN | 0.565371 | 0.6765916 | 0.1047328 | -0.190846 | -0.969264 | 146 |
| P53_PATHWAY | 0.5291829 | 0.6547517 | 0.11524 | -0.188616 | -0.984029 | 186 |
| XENOBIOTIC_METABOLISM | 0.4351145 | 0.5760776 | 0.1275053 | -0.196019 | -1.020512 | 180 |
| DNA_REPAIR | 0.3916084 | 0.5393851 | 0.1287887 | -0.202462 | -1.025112 | 143 |
| MITOTIC_SPINDLE | 0.3083004 | 0.4894318 | 0.1574029 | -0.201133 | -1.059736 | 196 |
| HEME_METABOLISM | 0.2382813 | 0.4460136 | 0.1797823 | -0.208979 | -1.083342 | 183 |
| PI3K_AKT_MTOR_SIGNALING | 0.2684564 | 0.4557515 | 0.155242 | -0.230794 | -1.089084 | 101 |
| ALLOGRAFT_REJECTION | 0.1535581 | 0.3296982 | 0.222056 | -0.217158 | -1.130669 | 177 |
| APOPTOSIS | 0.1152416 | 0.262895 | 0.2572065 | -0.230564 | -1.168771 | 151 |
| UV_RESPONSE_UP | 0.1104016 | 0.2599779 | 0.2878051 | -0.241811 | -1.224265 | 149 |
| INFLAMMATORY_RESPONSE | 0.0745861 | 0.2297331 | 0.2878051 | -0.237342 | -1.240252 | 188 |
| IL6_JAK_STAT3_SIGNALING | 0.0859873 | 0.2324841 | 0.2765006 | -0.274215 | -1.26288 | 85 |
| TNFA_SIGNALING_VIA_NFKB | 0.0127979 | 0.0849312 | 0.3807304 | -0.261052 | -1.367376 | 191 |
| FAvsAF.DN | 0.003455 | 0.028024 | 0.4317077 | -0.301439 | -1.523257 | 142 |
| UNFOLDED_PROTEIN_RESPONSE | 0.0006542 | 0.00796 | 0.4772708 | -0.350003 | -1.688202 | 108 |
| MTORC1_SIGNALING | 1.94E-05 | 0.000707 | 0.5756103 | -0.333941 | -1.749167 | 191 |
| MYC_TARGETS_V2 | 0.0019379 | 0.020209 | 0.4550599 | -0.410217 | -1.765031 | 58 |
| INTERFERON_ALPHA_RESPONSE | 2.92E-05 | 0.0007116 | 0.5756103 | -0.420069 | -1.954286 | 91 |
| INTERFERON_GAMMA_RESPONSE | 8.95E-08 | 6.53E-06 | 0.7049757 | -0.378937 | -1.980826 | 189 |

For the 175 samples with microarray expression AND "RESPONSE" data recorded, the samples were split by treatment-stage info: Pre-treatment (n=95) and Post-treatment (n=80). Then for each treatment-stage diff.-exp. was performed for "Non-responders Vs. Responders". Hence +ve enrichment score indicates enrichment in Non-responders.
