## Extended data figures for "Mutational order and epistasis determine the consequences of *FBXW7* mutations during colorectal cancer evolution"

### Extended data figure 1

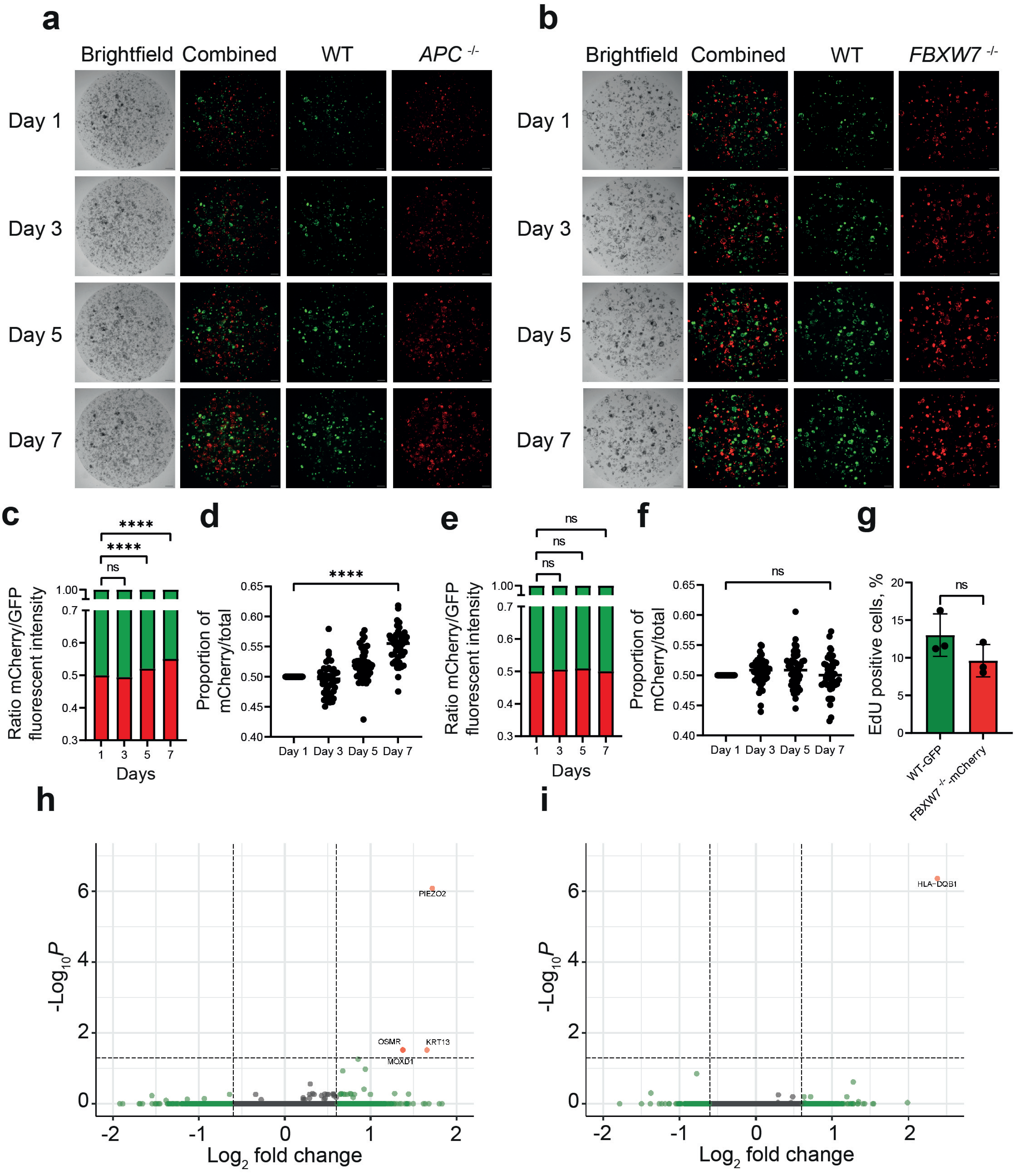


**Extended data figure 1 FBXW7-/- organoids do not influence growth of neighbouring wild-type organoids**

*APC*$^{-/-}$ cells act as supercompetitors which suppress the growth of neighbouring wildtype cells, evidenced by proliferative advantage of A organoids over neighbouring W organoids (**a, c, d**). A similar effect was not seen in F organoids co-cultured with W organoids (**b, e, f**). **g**, EdU assay for W and F organoids showed no significant differences in proliferation rate. Volcano plot of co-cultured W versus non-co-cultured W (**h**) and co-cultured F versus non-co-cultured F (**i**) organoids showed that the effect of co-culture did not significantly influence the transcriptome. Positive fold change represents upregulated in co-cultured. The values in this figure are mean ± sd, and statistical significance was measured by unpaired t-test. All experiments were performed with n=3 biological replicates.

### Extended data figure 2

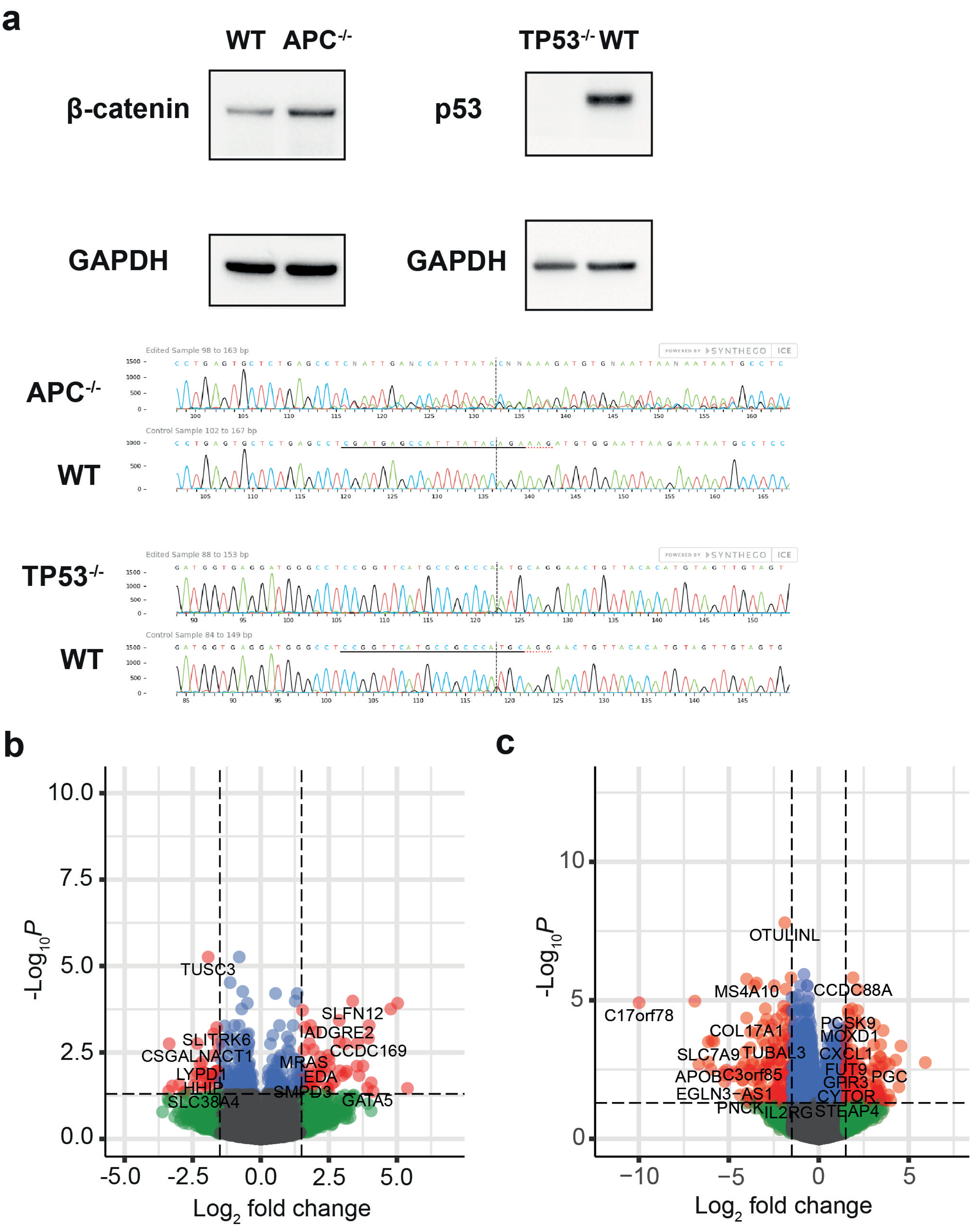


**Extended data figure 2 Mutational background influences the transcriptomic effect of an FBXW7 mutation**

**a**, Western blots for beta-catenin and p53 in wildtype and CRISPR-edited organoid. Sanger traces for edited and wildtype organoids as analysed by ICE tool (Synthego). **b**, RNAseq volcano plot comparing transcriptomes of AF vs A organoids. Positive fold change represents upregulated in AF. **c,** RNAseq volcano plot comparing transcriptomes of ATF vs AT organoids. Positive fold change represents upregulated in ATF. All experiments were performed with n=3 biological replicates.

### Extended data figure 3

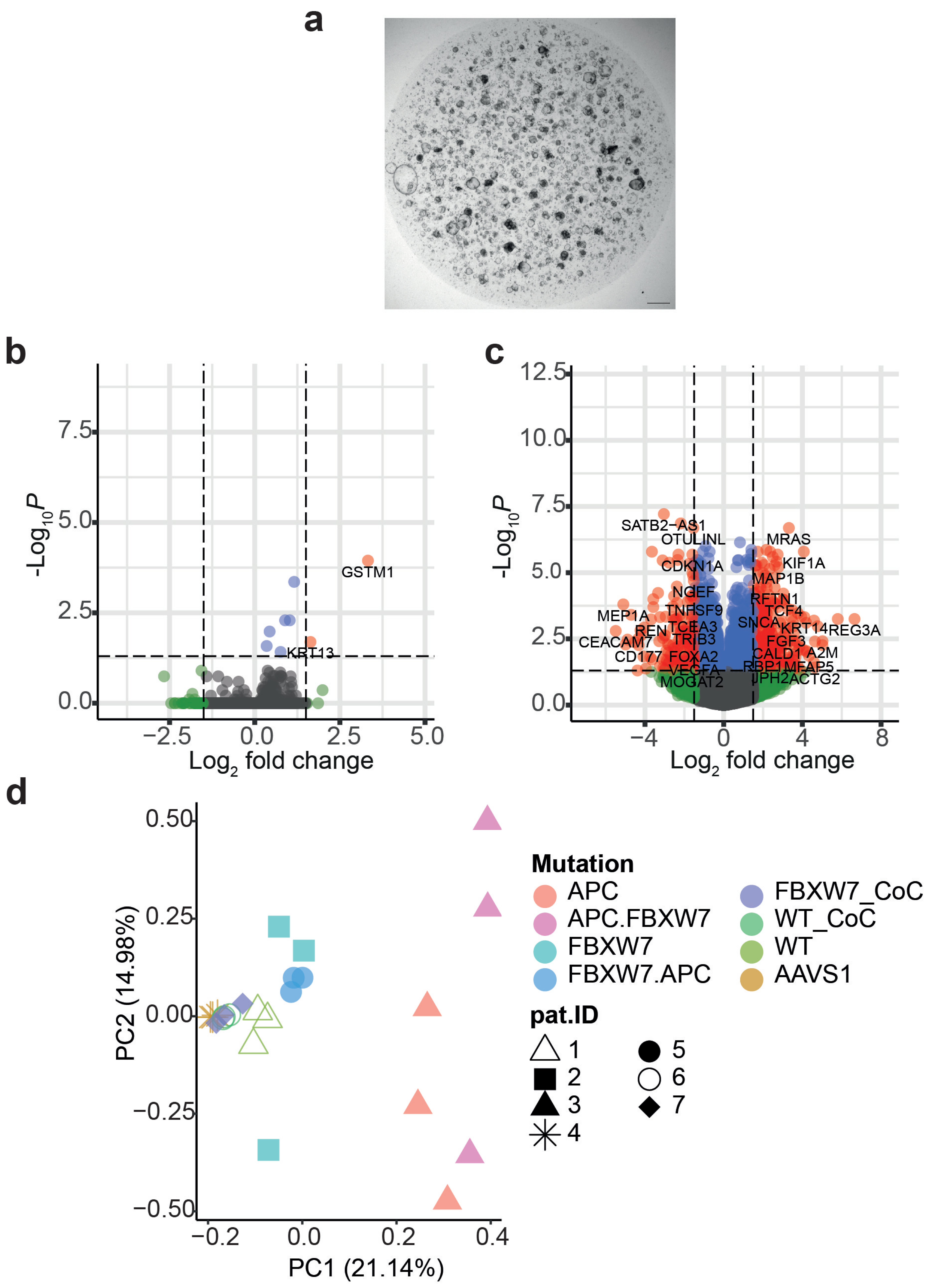


**Extended data figure 3 Effect of epistasis not an artefact of CRISPR/Cas9 gene editing**
**a**, Brightfield microscopy of organoids which had been gene edited at the AAVS1 loci three times to mimic triple-mutant ATF organoids. AAVS1 organoids did not adopt a cystic phenotype. Scale bar = 100um. **b**, RNAseq volcano plot comparing transcriptomes of AAVS1 vs W organoids. Positive fold change is upregulated in AAVS1 samples. **c**, RNAseq volcano plot comparing transcriptomes of ATF vs W organoids. Positive fold change is upregulated in ATF samples. All experiments were performed with n=3 biological replicates. d, Principal component analysis of transcriptomes from all gene-edited models identified by genotype and patient donor.

### Extended data figure 4

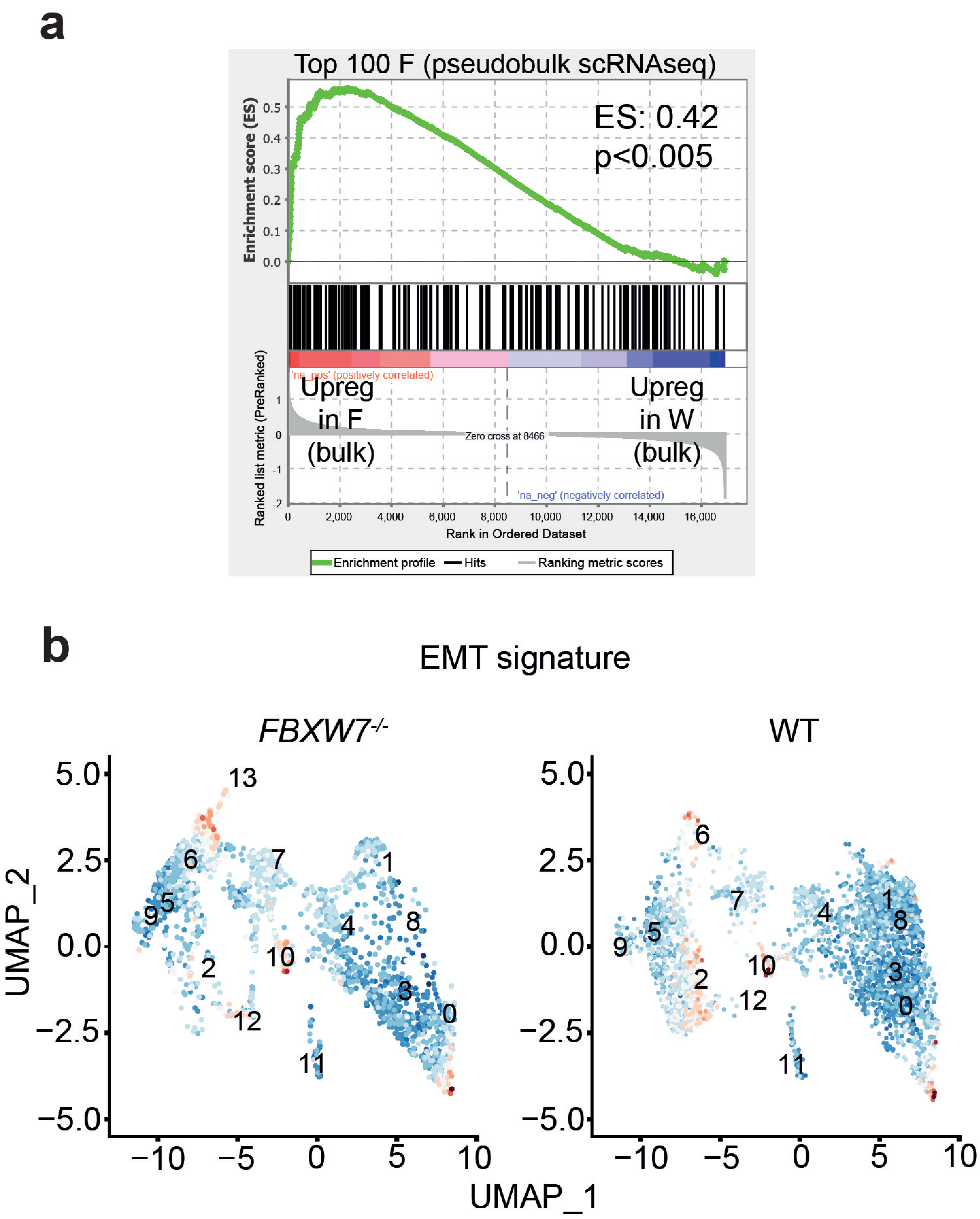


**Extended data figure 4 Cluster 13 in F organoids was marked by EMT signature**
**a**, GSEA plot of the enrichment of the top 100 F specific genes identified through pseudobulk analysis of scRNAseq data compared to ranked bulk RNAseq data derived from F vs W comparison. **b**, UMAP scRNAseq plots comparing F and W organoids with the EMT signature colour coded. Red = enriched. Blue = de-enriched

### Extended data figure 5

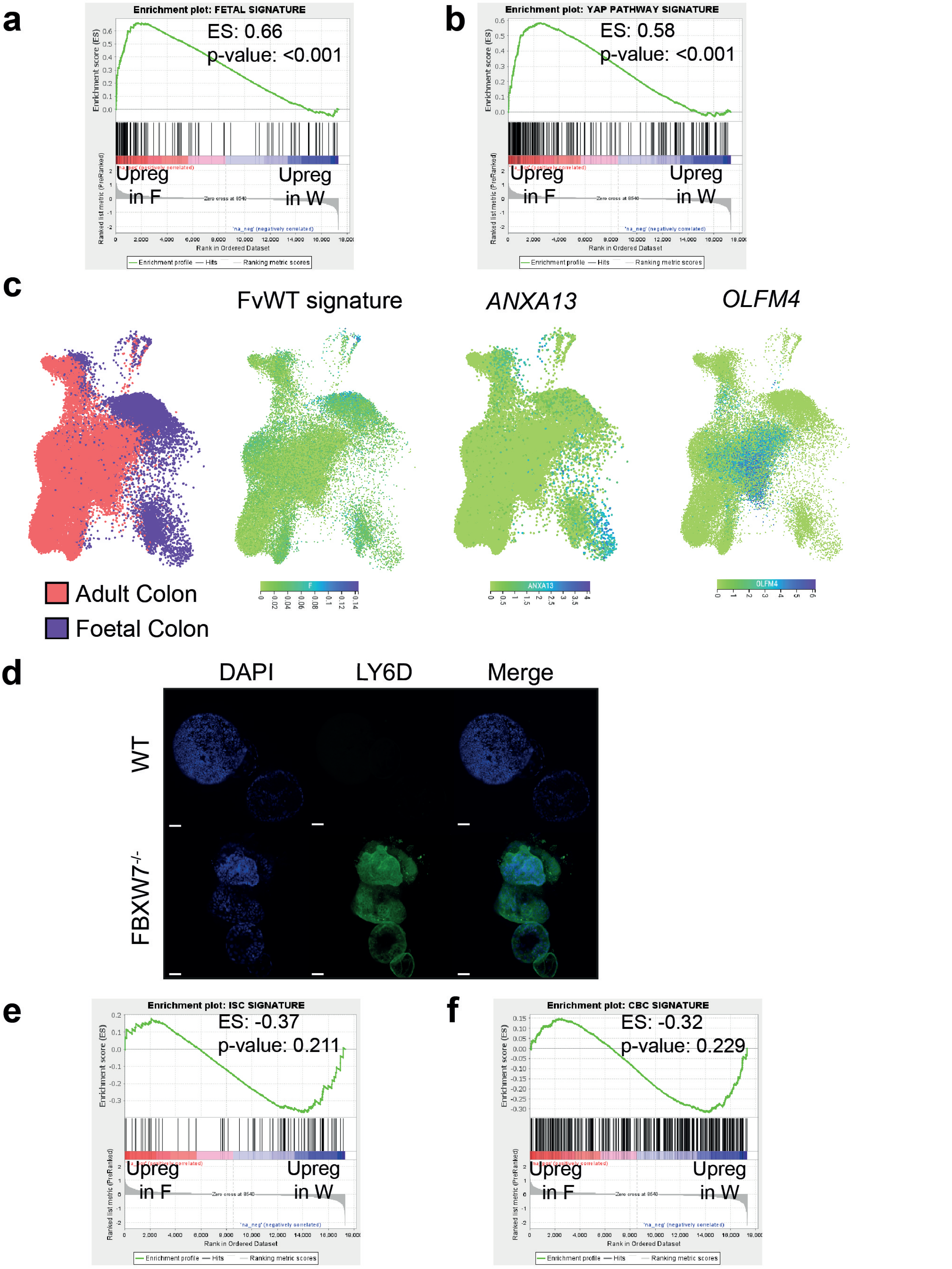


**Extended data figure 5 GSEA of bulk RNAseq data of F vs W organoids using key pathways derived from scRNAseq analysis**

GSEA plots for enrichment of the foetal signature [17] (**a**), and YAP pathway (**b**) in F organoids compared with W organoids. **c**, UMAP scRNAseq plots from Elmentaite et al of human, colon adult and foetal epithelial cells overlaid with enrichment for FvW signature, *ANXA13* and *OLFM4* expression. Blue = enriched. **d**, Immunofluorescence for LY6D in F and W organoids (Blue = DAPI, Green = anti-LY6D) Scale bars = 50um. GSEA plots for enrichments of the ISC (**e**) and CBC (**f**) signatures in F compared to W organoids.

### Extended data figure 6

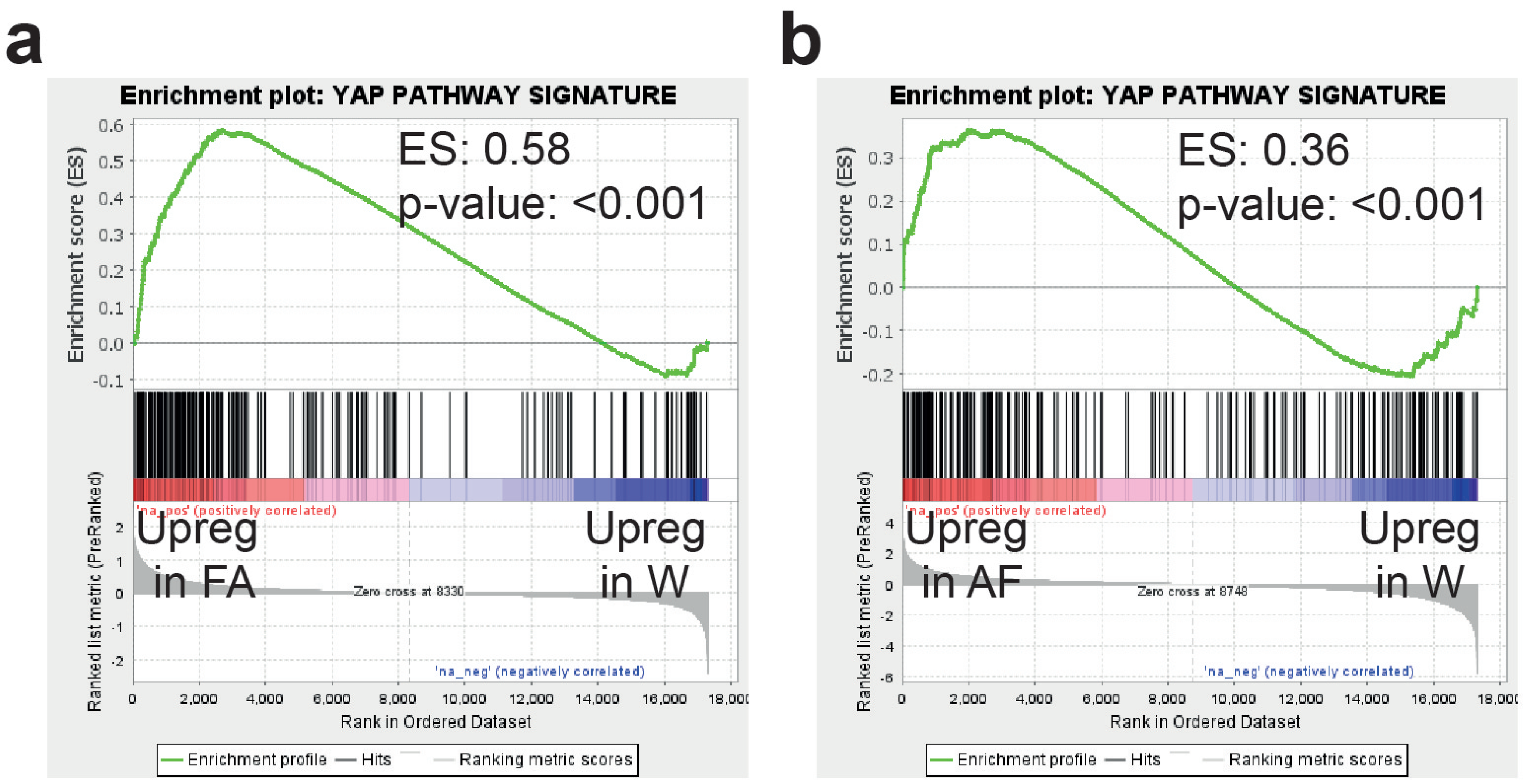


**Extended data figure 6 GSEA showing greater enrichment of YAP signalling in FA compared to AF organoids**
GSEA plots of bulk RNAseq data of FA vs W (**a**) and AF vs W (**b**) for the YAP pathway signature.

### Extended data figure 7

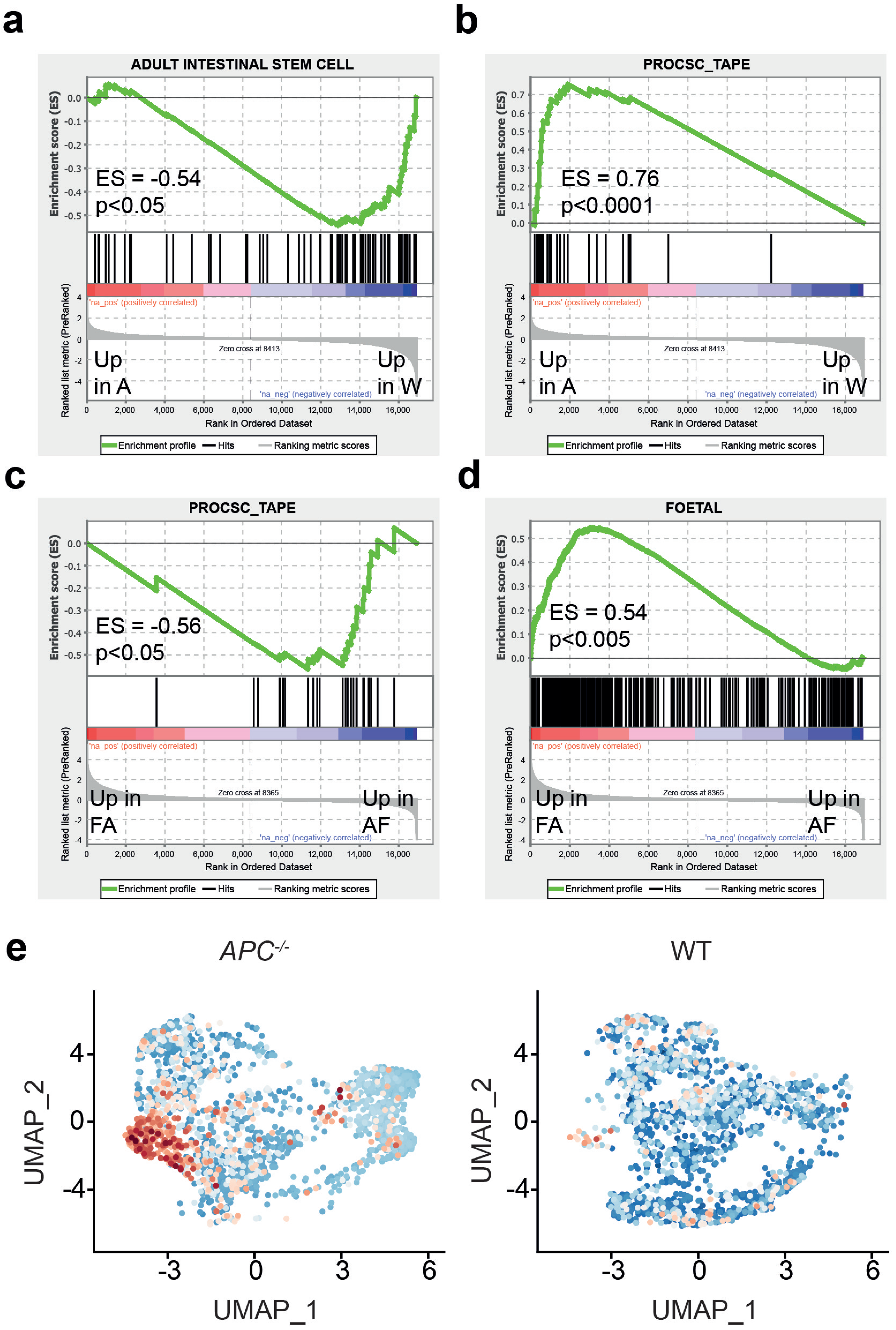


**Extended data figure 7 GSEA and scRNAseq analysis of A organoids compared to W**
GSEA plots of bulk RNAseq comparisons between A vs W for the adult intestinal stem cell signature (**a**) and proliferative cancer stem cell signature (**b**). GSEA plots of bulk RNAseq comparisons between FA vs AF for the proliferative cancer stem cell signature (**c**) and the foetal stem cell signature (**d**). **e**, UMAP single cell RNAseq plots of A and W organoids overlaid with enrichment for the proCSC signature (degree of red is equivalent to degree of enrichment).

### Extended data figure 8

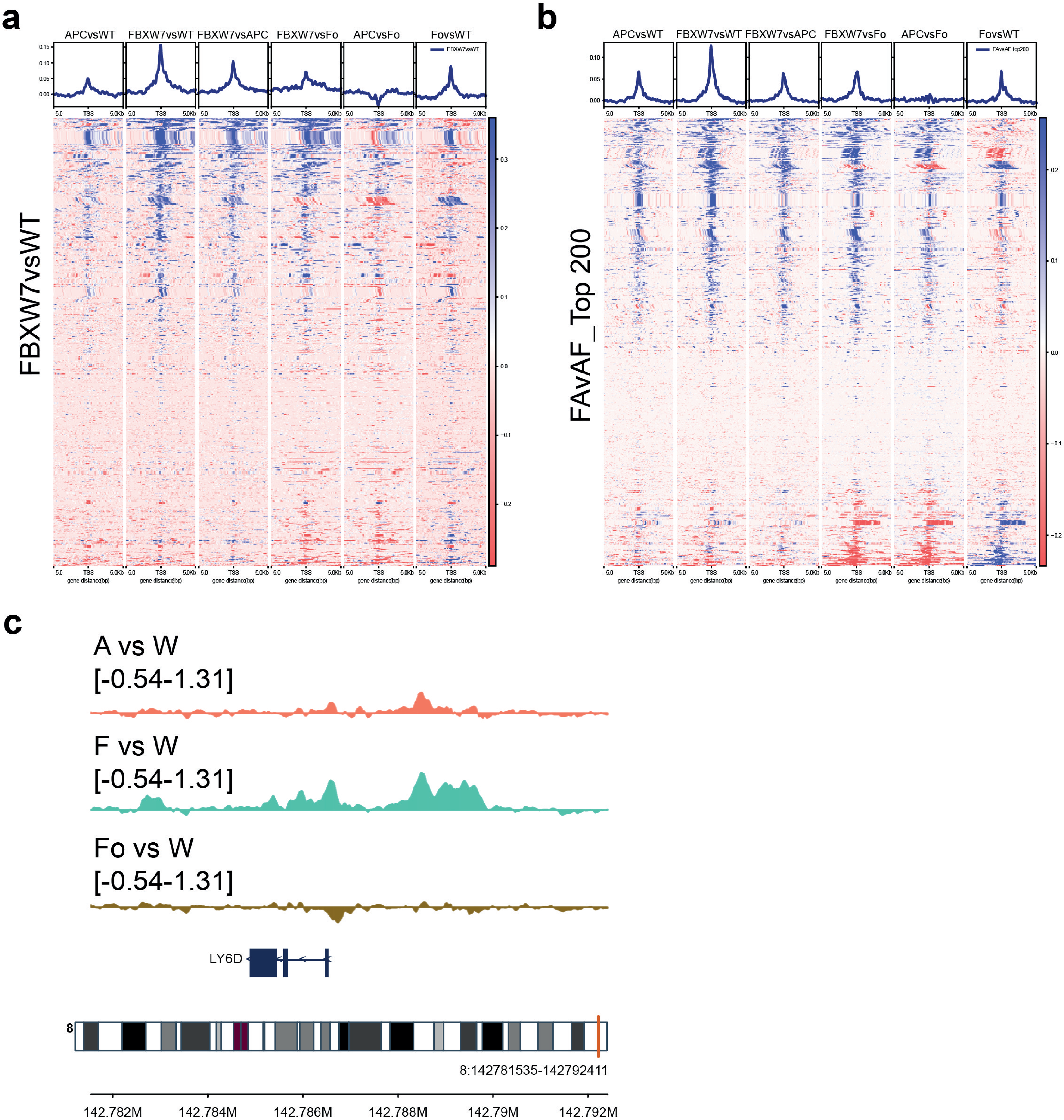


**Extended data figure 8 ATACseq profiles for accessibility of F vs W and FA vs AF signatures in W, A, F and Foetal organoids**

**a**, ATAC accessibility profiles for genomic loci associated with gene in the FvW RNAseq signature. (blue = increased accessibility, red = decreased accessibility). **b**, ATAC accessibility profiles for genomic loci associated with top 200 genes in the FavAF RNAseq comparison. (blue = increased accessibility, red = decreased accessibility). **c**, Chromatin accessibility for the *LY6D* gene region (chr8:142781535-142792411) in ‘A.vs.WT’, ‘F.vs.WT’ and ‘Foetal.vs.WT’ comparisons from ATAC-seq, visualised as track plots after group autoscaling of the input data. The Y-axis notes the track label and the range of the values within the region visualised. The vertical red bar on the ideogram of the chr8 indicates the location of the region. The X-axis indicates the direction of the genome, and the numbers (bottom-most track) are 2kb-spaced positions on the chr8. The canonical protein-coding transcript of LY6D gene (NM_003695) is represented as the dark-blue boxes (exons, n=3) connected by arrows (‘<’) on horizontal bar (introns).
